# Spatiotemporal molecular dynamics of the developing human thalamus

**DOI:** 10.1101/2023.08.21.554174

**Authors:** Chang N Kim, David Shin, Albert Wang, Tomasz J Nowakowski

**Affiliations:** Department of Neurological Surgery, University of California, San Francisco, CA, USA; Eli and Edythe Broad Center for Regeneration Medicine and Stem Cell Research, University of California, San Francisco, CA, USA; Department of Anatomy, University of California, San Francisco, CA, USA; Department of Psychiatry and Behavioral Sciences, University of California, San Francisco, CA, USA; Kavli Institute for Fundamental Neuroscience, University of California, San Francisco, CA, USA; Weill Institute for Neurosciences, University of California, San Francisco, CA, USA

## Abstract

The thalamus plays a central coordinating role in the brain. Thalamic neurons are organized into spatially-distinct nuclei, but the molecular architecture of thalamic development is poorly understood, especially in humans. To begin to delineate the molecular trajectories of cell fate specification and organization in the developing human thalamus, we used single cell and multiplexed spatial transcriptomics. Here we show that molecularly-defined thalamic neurons differentiate in the second trimester of human development, and that these neurons organize into spatially and molecularly distinct nuclei. We identify major subtypes of glutamatergic neuron subtypes that are differentially enriched in anatomically distinct nuclei. In addition, we identify six subtypes of GABAergic neurons that are shared and distinct across thalamic nuclei.

**One-Sentence Summary:** Single cell and spatial profiling of the developing thalamus in the first and second trimester yields molecular mechanisms of thalamic nuclei development.

## Main Text

Thalamus can be anatomically subdivided into multiple nuclei with distinct patterns of neuronal projections (*1*). Many thalamic subregions are hypothesized to have undergone extensive remodeling and reorganization in recent evolution, including the pulvinar nucleus that is extensively expanded in humans and primates compared to mice (*2, 3*). Thalamic neurons emerge from progenitor cells located in the diencephalon of the prenatally developing nervous system (*4*). Recent studies have begun to apply single cell genomics approaches to uncover the molecular dynamics of their differentiation in mice (*5*), but our understanding of thalamic neuron differentiation in humans is less well-characterized. To better understand their molecular and spatial organization, we investigated developing human thalamus during the first and second trimester using single cell and spatial transcriptomics.

### Neurogenesis in the first trimester thalamus

To identify the molecular trajectories of cell types in the developing human thalamus during neurogenesis, we performed an analysis of scRNA-seq datasets generated from samples from the first trimester of human development (*6, 7*). Across 5 samples (3 males and 2 females) between gestational week (GW) 6 to 10, data re-processing retained 27,362 cell transcriptomes passing quality control metrics generated using droplet-based scRNA-seq (10X Chromium v2 assay) (**fig. S1A**). Initial preprocessing steps include ambient RNA removal using FastCAR (*8*), doublet filtering with scds (*9*), and further cell filtering using spliced/unspliced ratio with DropletQC (*10*). We then performed SCTransform normalization (*11*) with Harmony (*12*) for batch correction between different samples and Louvain clustering to identify cell types (**Fig. 1, A and B**). We recovered ten clusters representing all of the major classes in the thalamus, including radial glia (RG1-2), intermediate progenitor cells (IPC1-2), GABAergic inhibitory neurons (IN1-3), glutamatergic excitatory neurons (EN1-2), and neural crest-derived mesenchymal stem cells (MES) that develop to form the cerebral vasculature (**Fig. 1B)**(*13*).

**Fig. 1.**
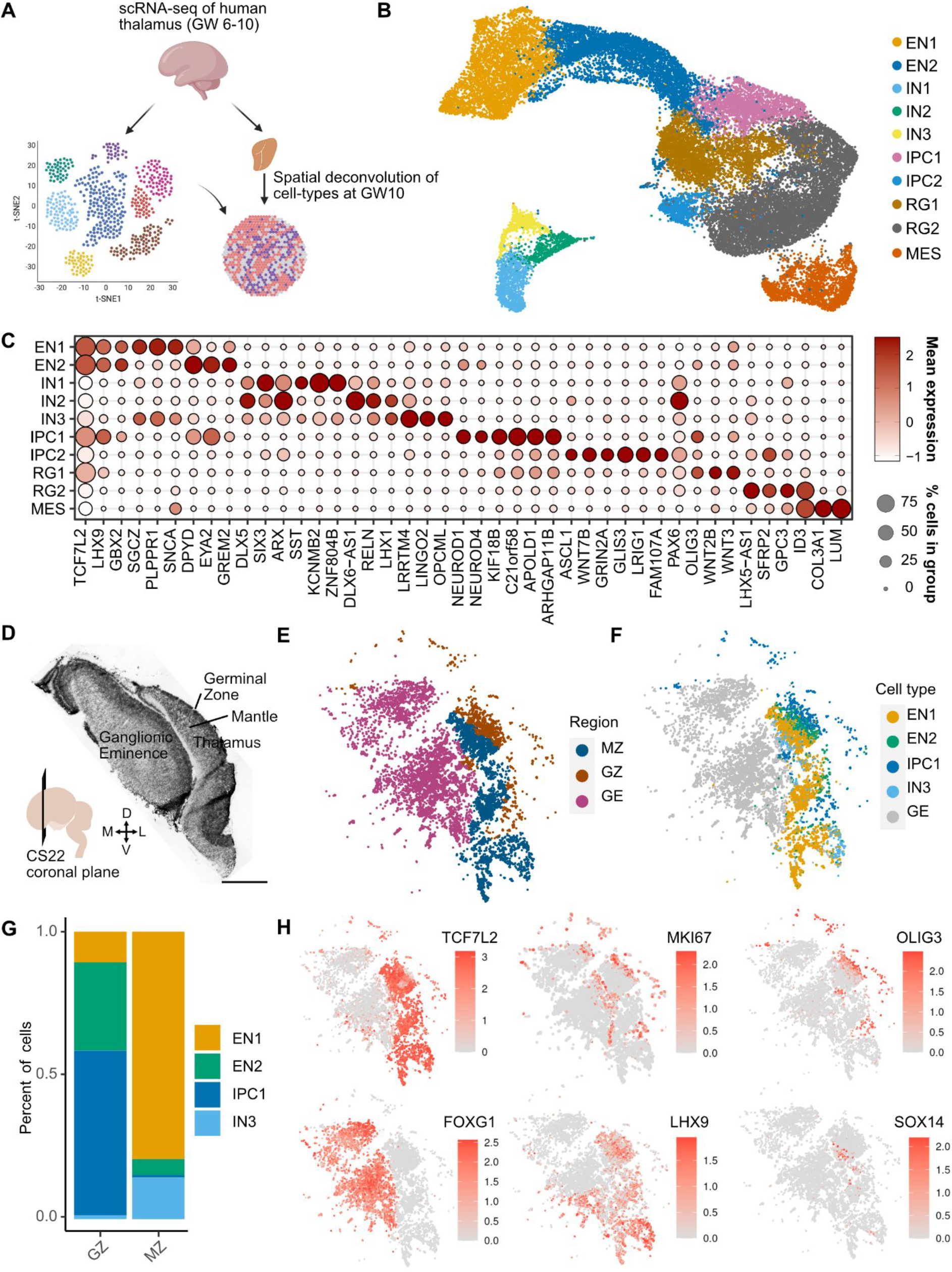
Neurogenesis in the first trimester thalamus. **A**. Schematic overview of figure 1. We analyzed scRNA-seq data collected from three biologically independent first trimester thalamus samples (GW6-10). Spatial transcriptomics using MERFISH was performed to spatially map cell types identified by scRNA-seq. **B**. UMAP embedding of cell type annotations on the Harmony batch corrected space: EN1/2 - excitatory neuron 1/2, IN1/2/3 - inhibitory neuron 1/2/3, RG1/2 - radial glia 1/2, IPC1/2 - intermediate progenitor cells 1/2, and MES - mesenchyme. **C**. Dot plot of marker genes across cell types/states. **D**. H&E of a MERFISH adjacent coronal section of CS22 (GW10) **E**. MERFISH region annotations. MZ - Thalamic Mantle Zone, GZ - Thalamic Germinal Zone, GE - Ganglionic Eminences. **F**. Spatially projected cell type annotations. **G**. Relative proportion of cell types within regions from E. **H**. Spatially projected expression levels of select marker genes, including telencephalic (*FOXG1*) and diencephalic (*TCF7L2, LHX9*) identity, dividing cells (*MKI67*), radial glia and IPCs (*OLIG3*), and rostral thalamus-derived GABAergic neurons (*SOX14*).

Radial glia are neural stem cells in the developing brain. In the embryonic diencephalon, radial glia line the third ventricle and are partitioned into three zones: prosomere 1, which gives rise to the pretectum, prosomere 2, which gives rise to the thalamus and epithalamus, and prosomere 3, which gives rise to the prethalamus (*14*). They express *PAX6* but lack the expression of IPC markers *NEUROD1, NEUROG1, NEUROG2*, and *ASCL1*, or neuronal markers such as *SLC17A6* and *GAD1 (15-17)*. We identified two clusters of radial glia (RG1-2) (**Fig. 1B**). RG1 are prosomere 2-derived thalamic radial glia based on their expression of *OLIG3*, a marker for radial glia and IPCs of thalamic glutamatergic lineage (**Fig. 1C**) (*18*). RG1 was enriched for genes associated with the Wnt pathway, including the Wnt ligands *WNT2B* and *WNT3*, the Wnt co-receptor *LRP6*, and a negative feedback regulator *ZNRF3*, consistent with the importance of Wnt signaling for thalamus development (**Fig. 1C and fig. S1D**) (*19, 20*). Other enriched genes included *GLI2*, a mediator of sonic hedgehog signaling, and the Rho GTPase *CPNE8* (**fig. S1D**). Despite the lack of canonical IPC marker expression, these cells also express *LHX9*, a marker associated with thalamic IPCs and glutamatergic neurons in mouse and zebrafish (**Fig. 1C**) (*17, 21*). RG2 are likely radial glia of GABAergic lineage with representation from the prosomere 3-derived prethalamus based on the expression of *FOXD1* and *LHX5* (*22*), and prosomere 2-derived rostral thalamus based on expression of *NKX2*.*2* and *OLIG2* (**Fig. 1C; fig. S1, C and D**) (*23-28*). Other distinguishing marker genes include *LHX5-AS1, ID3, SFRP2*, and *GPC3* (**Fig. 1C**). *ID3* was previously described as a pan-radial glial marker in mouse diencephalon including the thalamus (*29*), but *ID3* appears to be specific to RG2 and MES in humans (**Fig. 1C)**. A sparse population of *PAX3/LMO1*-expressing prosomere 1/pretectal progenitors (*5, 24, 30*) and *SHH/SIM2*-expressing zona limitans intrathalamica (ZLI) progenitors that reside between the thalamus and prethalamus (*5, 31, 32*) also cluster with RG2, suggesting that there was not enough representation from these populations to cluster separately (**fig. S1, C and D**). We also identified two IPC populations. IPC1 represents thalamic glutamatergic IPCs that express canonical markers that include *OLIG3, NEUROD1, NEUROG1, NEUROG2*, and *NEUROD4* (**Fig. 1C and fig. S1C**) (*17, 33*). We found additional novel markers for IPC1, including *KIF18B, C21ORF58, APOLD1, EYA2*, and *ARHGAP11B*, a human-specific gene that has been implicated in the evolutionary expansion of the human neocortex (**Fig. 1C**) (*34-38*). Preferential expression of *ARHGAP11B* in thalamic IPCs suggests that it may be involved in the elaboration of the human thalamus, similar to its known role in the cortex. IPCs of GABAergic lineage (IPC2) from prosomeres 2 and 3 express the GABAergic proneural transcription factor *ASCL1* (**Fig. 1C**) (*17*). This cluster was additionally enriched for the Wnt ligand *WNT7B*, the NMDA receptor subunit *GRIN2A*, the transcription factor *GLIS3*, the EGFR regulator *LRIG1*, and the tumor suppressor gene *FAM107A* (**Fig. 1C**).

Glutamatergic and GABAergic neurons were identified based on the expression of *SLC17A6* and *GAD1*, respectively. We identified two glutamatergic neuron populations in the scRNAseq dataset, EN1 and EN2, which pertain to thalamic neurons based on their expression of canonical markers *LHX9, LHX2*, and *GBX2* (**Fig. 1C and fig. S1D**) (*39, 40*). EN1 appear to be mature neurons based on their enrichment for *GAP43* (**fig. S1D**). These cells are additionally enriched for markers including *SGCZ, PLPPR1*, and *SNCA* (**Fig. 1C**). EN2 are newborn neurons based on co-expression of *GBX2*, a marker expressed in post-mitotic glutamatergic neurons in the thalamus (*39-41*) and IPC markers *NEUROD1* and *NEUROG2* (**Fig. 1C and fig. S1D**). These cells are enriched for *CACNA2D1, DPYD, KALRN*, and *EYA2* (**Fig. 1C and fig. S1D**). Three clusters represented GABAergic inhibitory neurons (IN1-3). IN1 and IN2 represent prosomere 3-derived prethalamic neurons, given their expression of canonical markers *PAX6, DLX5, SIX3*, and *ARX (16, 39, 42-44)* and absence of *FOXG1* expression, which is a pan-telencephalic marker (*45*) (**Fig. 1C and fig. S1D**). IN1 is additionally enriched for *ZNF804B, KCNMB2, ISL1*, and *SST*, a gene encoding for the neurotransmitter somatostatin (**Fig. 1C and fig. S1D**). IN2 is a *LHX1*-positive cluster enriched for *DLX6-AS1* and *RELN* (**Fig. 1C**). IN3 are likely prosomere 2/rostral thalamus (rTh)-derived GABAergic neurons, based on the expression of *LHX1, NXK2*.*2, OTX2, SOX14*, and genes encoding for the neurotransmitters neuropeptide Y (*NPY*) and enkephalin (*PENK*) (**fig. S1D)** (*18, 46-49*). Whereas *OTX2* and *SOX14* are also expressed in GABAergic neurons derived from midbrain and from prosomere 1-derived pretectum, *NKX2*.*2* expression is absent in pretectal and midbrain-derived GABAergic neurons (*18, 32, 50*). Furthermore, these cells lack expression of *LMO1* and *IRX3*, markers of caudal diencephalon also expressed in midbrain-derived GABAergic neurons, suggesting that the migratory stream of interneurons from the midbrain has not emerged during the first trimester (**fig. S1C and fig. S3D**) (*5, 51, 52*).

We performed MERFISH spatial transcriptomics on a coronal section from the GW10 rostral human forebrain to spatially identify cells from the glutamatergic thalamic lineage identified from scRNA-seq (**Fig. 1D**). We identified three distinct regions based on an H&E stain of an adjacent slice: the medial and lateral ganglionic eminences (GE), the thalamic germinal zone (GZ), and the thalamic mantle zone (MZ) (**Fig. 1D**). Our MERFISH analysis confirmed regional identity based on mutually exclusive expression of *TCF7L2* and *FOXG1* in the thalamus and GE, respectively (**Fig. 1H**). The GZ is enriched for *OLIG3*, and the MZ is enriched for *SLC17A6* (**Fig. 1H and fig. S1F**). *MKI67* expression segregated the ventricular zone (VZ) from the subventricular zone (SVZ) (**Fig. 1H)**. We performed clustering analysis of our MERFISH dataset in order to relate the scRNA-seq populations to the regional annotations (**Fig. 1, E and F**). We identified four clusters that corresponded to IPC1, EN1, EN2, and IN3 from our single cell sequencing dataset (**Fig. 1F)**. IPC1, which co-express *OLIG3* and *EYA2*, reside in the thalamic GZ (**Fig. 1, G and H; fig. S1F)**. Newborn neurons (EN2) expressing *GBX2* and *EYA2* are localized to the GZ and MZ, with bias for the GZ **(Fig. 1F-H; fig. S1F)**. Mature *LHX9/SLC17A6-*expressing glutamatergic neurons (EN1) are predominantly found in the MZ **(Fig. 1F-H; fig. S1F)**. We also observed the presence of *SOX14/NKX2*.*2*-expressing GABAergic neurons (IN2) associated with rostral thalamus origin in the MZ **(Fig. 1F-H; fig. S1F)**. *PAX6/OLIG3*-expressing radial glia are observed at the VZ (**Fig. 1H and fig. S1F)**. However, due to damage at the VZ, there were likely too few of these cells in our MERFISH data to form a discrete cluster.

Collectively, our analysis uncovers the cellular diversity of the human thalamus and their spatial distribution during the first trimester. We identify *OLIG3*-positive thalamic radial glia and IPCs, and the glutamatergic neurons that they likely give rise to that classify broadly into two molecular subtypes. In addition, we define *ID3/ASCL1*-expressing neural progenitors, and their presumed GABAergic neurons progeny, which can be classified into three subtypes of diencephalon-derived GABAergic neurons. These cell types express similar patterns of marker genes compared to what has been previously observed in other species, in addition to genes not previously associated with thalamic cell type-enriched expression. These findings suggest that the overall molecular blueprint of first trimester diencephalic cell types is conserved in humans. The majority of GABAergic neurons in the thalamus has been reported to originate from extra-thalamic regions, including the midbrain in mice (*50, 53*) and possibly the ganglionic eminences in humans (*54, 55*). However, we do not observe evidence for their presence in the first trimester, suggesting that these migratory streams develop later. Spatially, the thalamus has not yet partitioned into nuclei, based on cytoarchitecture revealed by Nissl staining or by molecular profiles revealed by MERFISH **(Fig. 1D-F)** (*56*). As expected for this stage of development, thalamic cells segregate into layers that relate to the germinal zones containing radial glia and intermediate progenitor cells, and the mantle zone containing migrating and maturing neurons.

### Differentiation of thalamic neurons in the second trimester

Next, to gain insight into the molecular mechanisms and cellular composition of early thalamic nuclei, we analyzed scRNA-seq and snRNA-seq datasets from the second trimester of development, which were derived from 10 specimens (4 female and 6 male) between GW16 and GW25. This analysis retained 137,761 high quality cells or nuclei. For a subset of the samples, metadata information for thalamic subdivisions had been retained from dissections (*6*) based on microdissections along the dorsal-ventral and rostral-caudal axes, and included the pulvinar nucleus (**fig. S4A**). We identified 23 cell clusters, including all major cell classes: glutamatergic neurons (EN1-2), GABAergic neurons (IN1-6), intermediate progenitor cells (IPC), astrocytes (AC1-2), glial progenitor cells (GL), dividing cells (DIV), microglia (MG), ependymal cells (EP), oligodendrocyte progenitor cells (OPC), and oligodendrocytes (OL) (**Fig. 2A**). We also identified several vascular cell classes including endothelial cells (EC), pericytes (PC), and fibroblasts (FB) that were annotated based on our recent transcriptomic atlas of the human cerebrovasculature (**Fig. 2A**) (*23*).

**Fig. 2.**
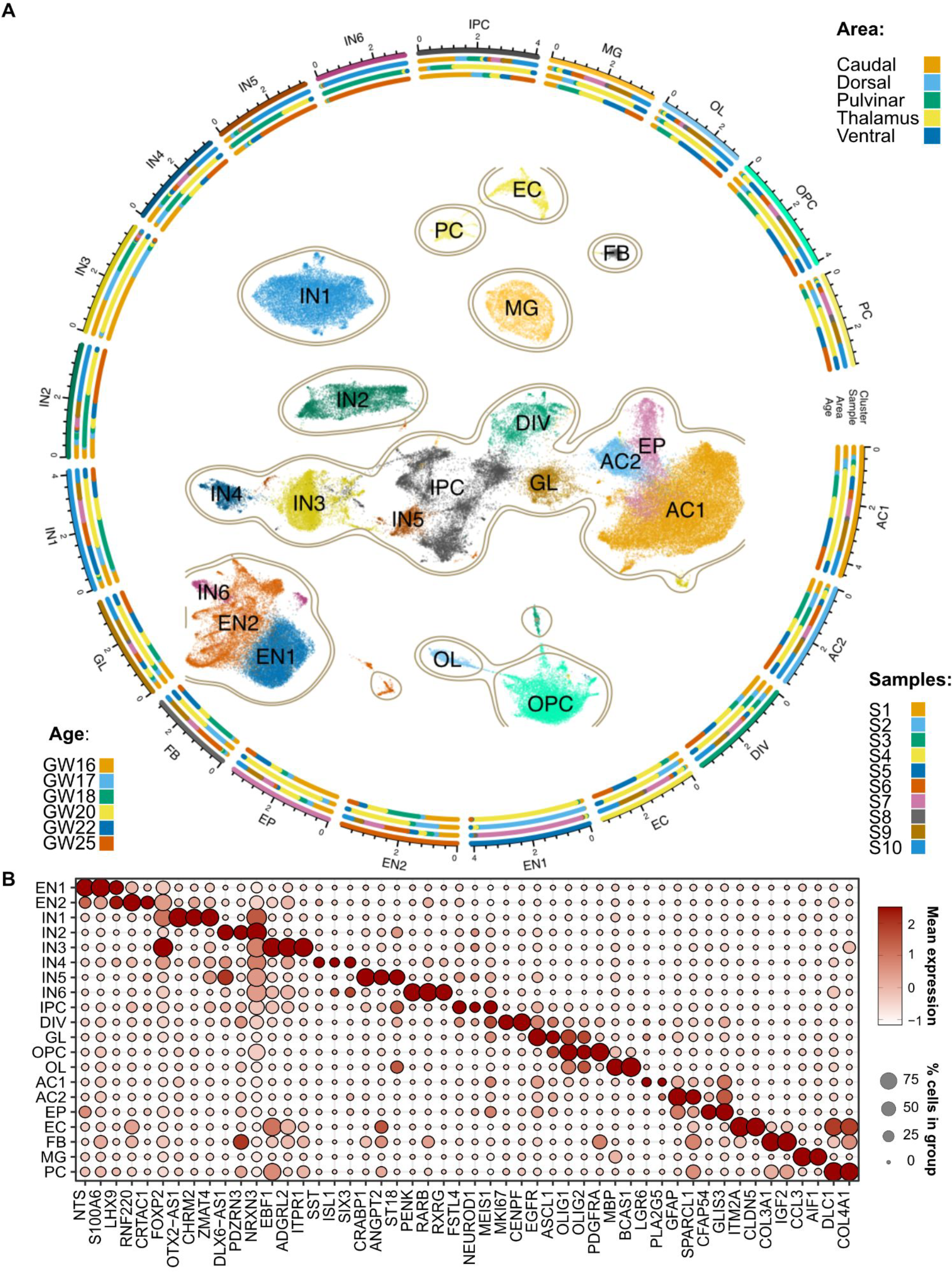
Cellular diversity in the second trimester thalamus. **A**. Constellation plot representing cellular diversity in second trimester human thalamus. PC - pericytes, OPC - oligodendrocyte precursor cells, OL - oligodendrocytes, MG - microglia, IPC - intermediate progenitor cells, IN1-6 - Inhibitory GABAergic neurons 1-6, GL - glial progenitor cells, FB - fibroblasts, EP - ependymal cells, EN1-2 - excitatory glutamatergic neurons 1-2, EC - endothelial cells, DIV - dividing cells, AC1-2 - astrocytes 1-2. **B**. Dot plot represents enrichment of select marker gene expression across cell types in the second trimester human thalamus.

Neural and glial progenitors were identified based on their expression of *HES1* **(fig. S3D)**. However, unlike in first trimester, we did not detect *OLIG3*+ clusters associated with thalamic progenitor cells **(fig. S3D)**, in line with the early window of neurogenesis observed in mouse (*57-59*). However, we identified *MKI67*+ dividing cells (DIV) and *NEUROD1/LHX1*+ IPCs (IPC) that were enriched for *MEIS1* and *PAX3*, markers of pretectal progenitors **(Fig. 2B and fig. S3D) (*5*)**. This observation suggests ongoing neurogenesis in the pretectum, consistent with prior findings in the diencephalon (*58*). We identified two molecularly distinct astrocyte populations, AC1 and AC2, based on their expression of known astrocyte markers *AQP4* and *GJA1 (60-63)* **(fig. S3D)**. These populations are differentiated by *LGR6* (AC1), a marker recently found to be enriched in thalamic astrocytes in the adult brain (*64*), and *SPARCL1* (AC2) **(Fig. 2B) (*65*)**. Ependymal cells (EP) exhibited similar transcriptomic profiles to astrocytes but were distinguished based on expression of *FOXJ1* **(fig. S3D)** *(66)*. We additionally observed a glial population (GL) that expresses *EGFR, ASCL1, OLIG1*, and *OLIG2*, but not *PDGFRA*, which is a signature of multipotent glial progenitor cells that may produce both astrocytes and oligodendrocytes **(Fig. 2B)** (*67, 68*). In contrast, OPCs and oligodendrocytes (OL) were distinguished from GL based on the expression of canonical markers *PDGFRA* and *MBP*, respectively **(Fig. 2B)** (*69, 70*). These results suggest that the end of neurogenesis and the onset of gliogenesis in the human thalamus occurs by GW16, which represents the earliest sample in our second trimester dataset.

Glutamatergic excitatory neuron clusters were defined based on shared expression of *LHX9* and *SLC17A6* **(Fig. 2B and fig. S3D)**, and consisted of two clusters (EN1-2). EN1 was enriched for neurotensin (encoded by *NTS)*, calcyclin (encoded by *S100A6*), and *SOX2*, whereas EN2 was enriched for *RNF220, CRTAC1*, and *FOXP2* **(Figs. 2B and 3B)**. EN1 and EN2 likely relate to the *SOX2* and *FOXP2-*expressing subtypes recently observed in the embryonic mouse thalamus (*5*). Three classification schemes exist for thalamic glutamatergic neurons in the adult brain for which molecular markers have been identified. The core/matrix model classifies thalamic neurons based on whether they project to layer IV or layer I in the cortex, and in primates can be molecularly identified based on their expression of *CALB1* or *PVALB* (*71*). The first order/higher order model relates to thalamic nuclei and their involvement in sensory relay pathways or transthalamic cortico-cortical communication (*72*), and their transcriptional signatures have been ascertained in mice (*73*). Finally, transcriptomic profiling of the different thalamocortical projection systems in mice revealed three major genetic profiles (primary, secondary, tertiary) found across thalamic pathways, and can be identified based on expression of *TNNT1, NECAB1*, or *CALB2* (*74*). We sought to determine whether any of these classification schemes could be applied to the second trimester human brain based on gene expression. We first explored whether EN1 and EN2 could be associated with core or matrix identity based on expression of *PVALB* and *CALB1*, respectively. These genes were detected in only a sparse number of cells, suggesting that these genes turn on later in development **(fig. S3D)**. We next sought to determine whether EN1 and EN2 were associated with transcriptional signatures of first order and higher order nuclei or with primary/secondary/tertiary profiles by performing gene signature scoring using the UCell package (*75*). We identified that EN1 was enriched for signatures of first order nuclei compared to EN2, but were not enriched for any of the primary/secondary/tertiary profiles **(Fig. 3C)**. However, EN1 was enriched for *CALB2*, a marker for tertiary profile neurons that relate to matrix neurons in intralaminar nuclei **(fig. S3D)** (*74*). In contrast, EN2 was enriched for signatures of higher order nuclei and secondary profile neurons compared to EN1, including *NECAB1* **(Fig. 3C and fig. S3D)**. Secondary profile neurons relate to matrix neurons that are enriched in higher order nuclei (*74*). Collectively, our data suggests that the genetic programs underlying sensory and higher-order relays may begin to be established during prenatal development in humans, prior to the onset of sensory experience. EN1 may be enriched for neurons associated with first order nuclei, and with matrix neurons in intralaminar nuclei based on expression of *CALB2*. EN2 is associated with matrix neurons enriched in higher order nuclei.

**Fig. 3.**
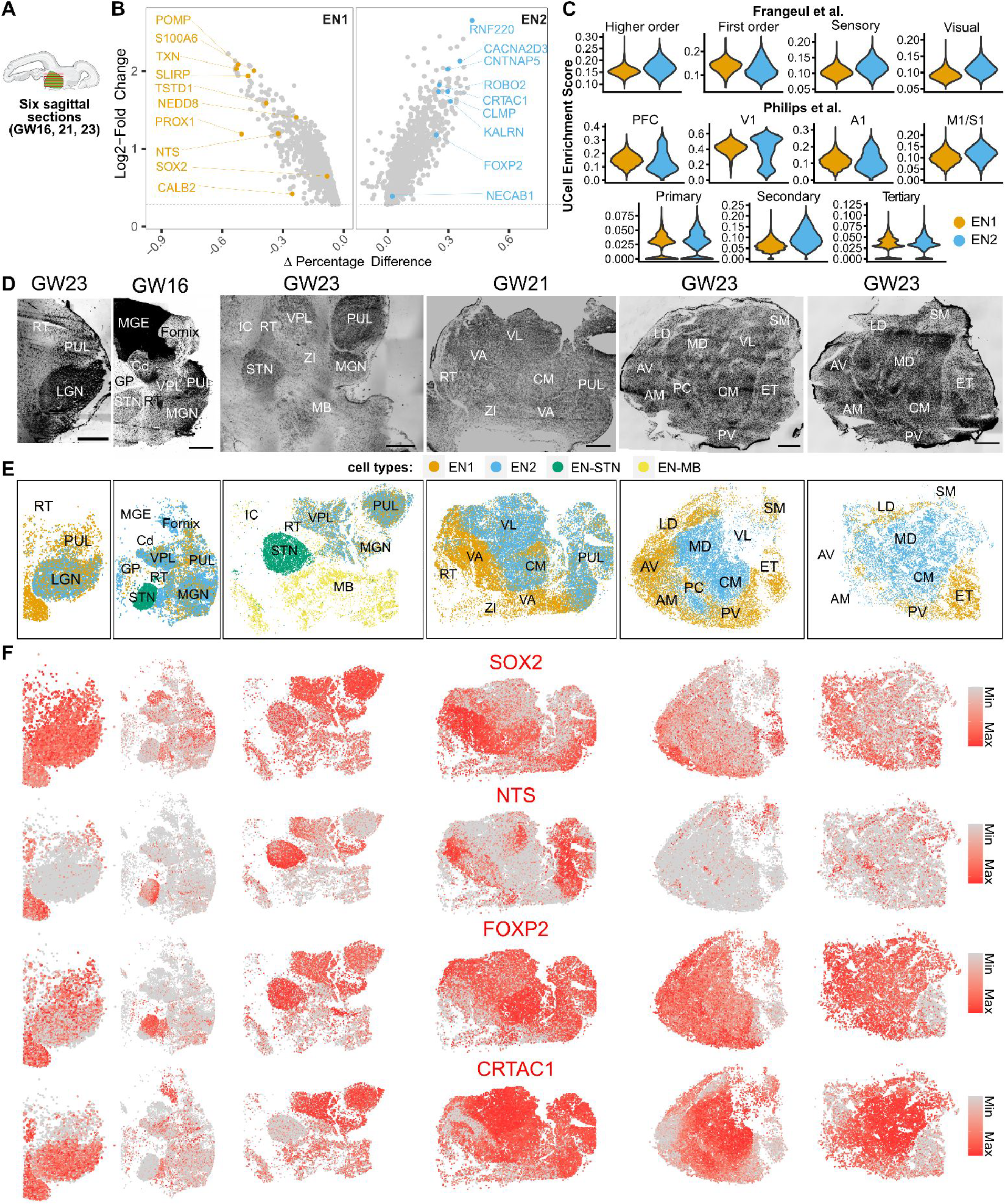
Spatially mapped thalamic glutamatergic neuron subtypes in human mid-gestation thalamus. **A**. Eight sagittal sections total were collected from three biologically independent second trimester thalamus specimens (GW16, 21, 23). Six representative sections are shown here and the other two are shown in **figs. S6 and S7. B**. Marker genes enriched for glutamatergic subtypes EN1 and EN2. **C**. UCell enrichment scores for gene signatures relating to known classes of thalamus neurons from adult mouse studies calculated for glutamatergic neuron subtypes EN1 and EN2. PFC - prefrontal cortex, V1 - Visual cortex, A1 - Auditory cortex, M1/S1 Motor/Somatosensory cortex. **D**. Nissl stains of sagittal sections reveal boundaries of thalamic nuclei. Sections are in order from left (lateral-most) to right (medial-most). Scale bar: 2 mm. RT - Reticular Nucleus, PUL - Pulvinar, LGN - Lateral Geniculate Nucleus, MGE - Medial Ganglionic Eminence, Cd - Caudate, GP - Globus Pallidus, STN - Subthalamic Nucleus, VPL - Ventral Posterior Lateral Nucleus, MGN - Medial Geniculate Nucleus, MB - Midbrain, VA - Ventral Anterior Nucleus, VL - Ventral Lateral Nucleus, CM - Centromedian Nucleus, ZI - Zona Incerta, AV - Anteroventral Nucleus, AM - Anteromedial Nucleus, LD - Dorsolateral Nucleus, MD - Dorsomedial Nucleus, PC - Paracentral Nucleus, PV - Paraventricular Nucleus, SM - Stria Medullaris, ET - Epithalamus. **E**. Spatial clustering analysis of MERFISH transcriptomics data reveals glutamatergic neuron subtypes. EN1 and EN2 from spatial clustering analysis relate to EN1 and EN2 from scRNA-seq data based on gene expression profiles. EN-STN and EN-MB are distinct clusters not represented by our scRNA-seq, and relate to glutamatergic neurons in the subthalamic nucleus and midbrain, respectively. **F**. Spatial feature plots reveal the expression of two example marker genes for EN1 (*SOX2, NTS)* and EN2 (*FOXP2, CRTAC1*) glutamatergic neuron subtypes in MERFISH datasets.

We also identified six clusters corresponding to GABAergic neurons based on the expression of *GAD1* and *SLC32A1* **(fig. S3D)**. IN1 expressed *OTX2* and *SOX14* but were mostly negative for *NKX2*.*2*, a marker for prosomere 2-derived GABAergic neurons, suggesting that they represent midbrain-derived GABAergic neurons **(fig. S3D)** (*53*). These cells were additionally enriched for *OTX2-AS1, CHRM2, ZMAT4*, and *LMO1* (**Fig. 2B and fig. S3D)**. Because this GABAergic neuron subtype was not detected in our first trimester data, we suspect that this population likely represents a migratory stream that enters the thalamus after GW10. IN3 expressed *EBF1, EBF3, ISLR2*, and *ESRRB*, suggestive of pretectal identity **(Fig. 2B and fig. S3D)** (*29, 76*). IN4 expressed markers of thalamic reticular nucleus including *SIX3, ISL1*, and *SST* **(Fig. 2B) (*77*)**. The remaining three populations (IN2, IN5, and IN6) expressed the pan-telencephalic marker *FOXG1* **(fig. S3D) (*45*)**. Previous anatomical studies have suggested that in humans, ganglionic eminence-derived GABAergic neurons migrate into thalamus (*26, 27*). *FOXG1*+ clusters could be distinguished based on differential enrichment for *PDZRN3* in IN2, *CRABP1, ANGPT2* and *MAF* in IN5, and *PENK, RARB*, and *RXRG* in IN6 (**Fig. 2A and fig. S3D**). IN5 is reminiscent of a recently described primate-specific *CRABP1/MAF1* positive interneuron population derived from the medial ganglionic eminences (MGE) (*28*). In summary, we identified GABAergic populations of diencephalic, midbrain, and telencephalic origin, the latter two of which emerge during the second trimester of development in humans.

### Spatial registration of thalamic cell type diversity

To investigate the spatial distribution of cell types in the developing human thalamus, we performed MERFISH on eight sagittal sections total collected from three biologically independent mid-gestation thalamus specimens using the Vizgen platform (**Fig. 3A**). These sections collectively captured the majority of thalamic nuclei as annotated by the Bayer and Altman atlas of the second trimester human brain (**Fig. 3D**) (*78*). Our gene panel was designed using cluster markers from second trimester sc/snRNA-seq data, which allowed us to visualize glutamatergic and GABAergic neurons, astrocytes, oligodendrocytes, microglia, endothelial cells, pericytes, and fibroblasts **(Table S3,Figs. 3, 4, and 5)**.

**Fig. 4.**
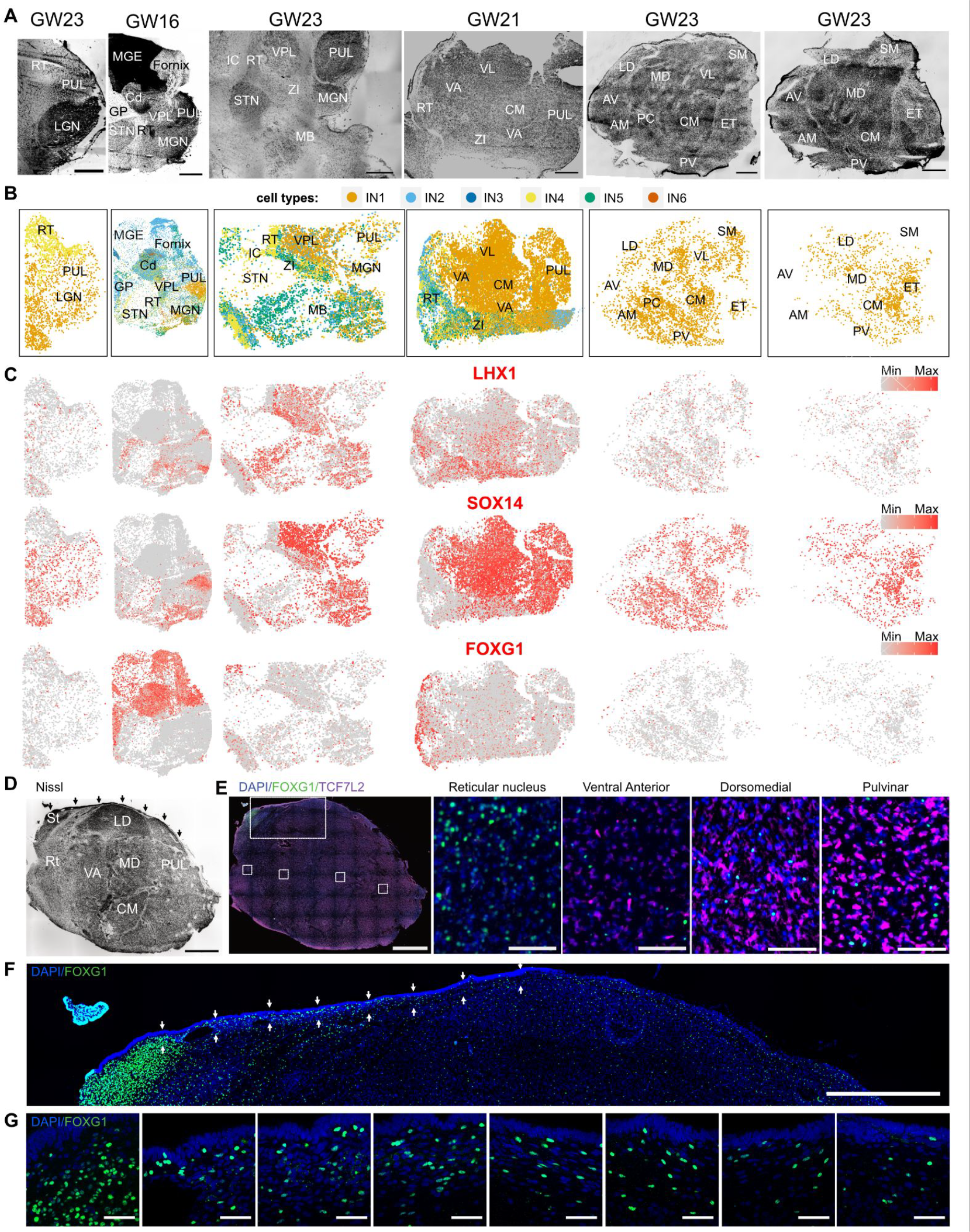
Spatially mapped GABAergic neuron subtypes across the human mid-gestation thalamus. **A**. Nissl stains of sagittal sections reveal boundaries of thalamic nuclei. Sections are in order from left (lateral-most) to right (medial-most). Scale bar: 2 mm. RT – Reticular Nucleus, PUL - Pulvinar, LGN - Lateral Geniculate Nucleus, MGE - Medial Ganglionic Eminence, Cd - Caudate, GP - Globus Pallidus, STN - Subthalamic Nucleus, VPL - Ventral Posterior Lateral Nucleus, MGN - Medial Geniculate Nucleus, MB - Midbrain, VA - Ventral Anterior Nucleus, VL - Ventral Lateral Nucleus, CM - Centromedian Nucleus, ZI - Zona Incerta, AV - Anteroventral Nucleus, AM - Anteromedial Nucleus, LD - Dorsolateral Nucleus, MD - Dorsomedial Nucleus, PC - Paracentral Nucleus, PV - Paraventricular Nucleus, SM - Stria Medullaris, ET - Epithalamus. **B**. Spatial clustering analysis of MERFISH transcriptomics data reveals GABAergic neuron subtypes IN1-6, which correspond to IN1-6 clusters from scRNA-seq data based on gene expression profiles. **C**. Expression of GABAergic neuron subtype markers across the sections. **D-E**. Nissl (D) and Immunostaining (E) of FOXG1 and TCF7L2 on a lateral section in a sagittal section from a biologically independent GW19 thalamus. Arrows in Nissl stain highlight the corpus gangliothalamicus, which is a thin migratory stream from MGE to thalamus. Small dotted boxes are zoom-in images depicted on the right. Large dotted box is a zoom-in image depicted in **Fig. 4F**. Scale bar for Nissl and low magnification images: 2 mm. Scale bar for high magnification images: 50 μm. St: Stria terminalis. **F**. Zoom in of the dorsal region of the thalamus section depicted in (E) to highlight FOXG1 staining within and below the corpus gangliothalamicus (highlighted by the arrows). Scale bar: 2 mm. **G**. High magnification (63X) images of regions highlighted by arrows in **Fig. 4F** highlights FOXG1 staining within the corpus gangliothalamicus, in order from left to right. Scale bar: 50 μm.

**Fig. 5.**
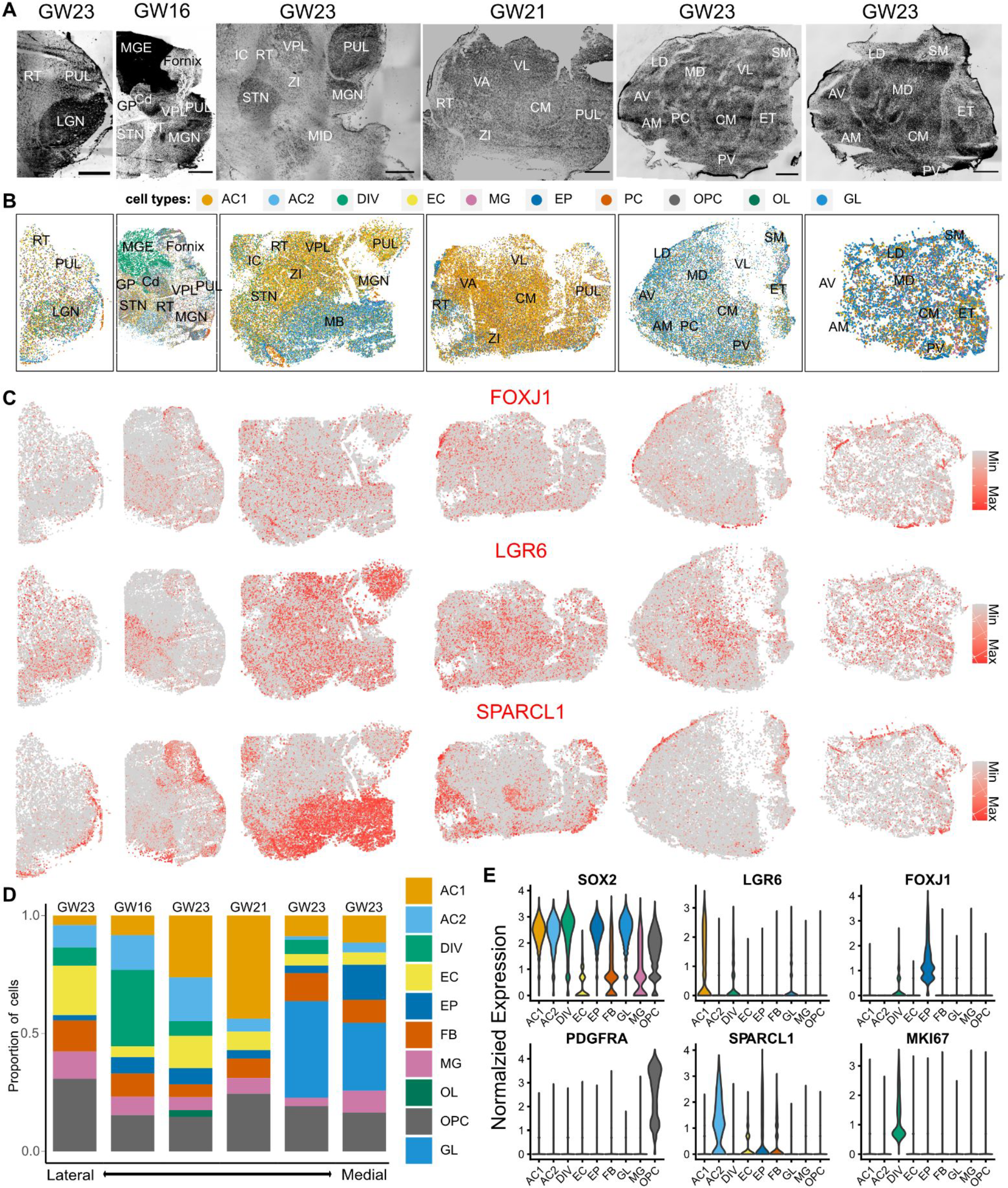
Spatially mapped non-neural cell subtypes across the human mid-gestation thalamus. **A**. Nissl stains of sagittal sections reveal boundaries of thalamic nuclei. Sections are in order from left (lateral-most) to right (medial-most). Scale bar: 2 mm. RT - Reticular Nucleus, PUL - Pulvinar, LGN - Lateral Geniculate Nucleus, MGE - Medial Ganglionic Eminence, Cd - Caudate, GP - Globus Pallidus, STN - Subthalamic Nucleus, VPL – Ventral Posterior Lateral Nucleus, MGN - Medial Geniculate Nucleus, MB - Midbrain, VA - Ventral Anterior Nucleus, VL - Ventral Lateral Nucleus, CM - Centromedian Nucleus, ZI - Zona Incerta, AV - Anteroventral Nucleus, AM - Anteromedial Nucleus, LD - Dorsolateral Nucleus, MD - Dorsomedial Nucleus, PC - Paracentral Nucleus, PV - Paraventricular Nucleus, SM - Stria Medullaris, ET - Epithalamus. **B**. Spatial clustering analysis of MERFISH transcriptomics data reveals non-neuronal subtypes. AC1 and AC2 - astrocyte subtypes, DIV - dividing cells, EC - endothelial cells, EP - ependymal cells, FB - fibroblasts, MG - microglia, OL - oligodendrocytes, OPC - oligodendrocyte progenitor cells, GL - bipotent glial progenitor cells. **C**. Spatial feature plots depicting normalized expression of EP (*FOXJ1*), AC1 (*LGR6*), and AC2 (*SPARCL1*) markers. **D**. Relative proportion of non-neural cell types across lateral-to-medial sections. **E**. Violin plots depicting normalized expression of glial, astrocytes, ependymal, and oligodendrocyte precursor marker genes across the 2 lateral-most GW23 sections.

We first analyzed the distribution of glutamatergic clusters EN1 and EN2 **(Fig. 3E; fig. S5B; fig. S7B; fig. S8; fig. S9)**. We observed an enrichment for EN1 neurons in rostral and medial regions of the thalamus, with a greater abundance of these cells observed in AV/AM, LD, VA, PC, and PV **(Fig. 3E)**. In contrast, EN2 exhibited a caudal-lateral bias, with an enriched distribution in the MD, VL, and CM **(Fig. 3E)**. Sensory nuclei such as LGN, MGN, and VPL contained similar abundance of both EN1 and EN2 populations **(Fig. 3E and fig. S7B)**. Pulvinar, which represents an evolutionarily expanded thalamic nucleus (*79*), shows an overall enrichment for EN1 in the lateral-most section and an enrichment for EN2 in more medial sections **(Fig. 3E)**. This difference within the pulvinar may relate to its functional organization, whereby lateral pulvinar is involved with the visual pathway and medial pulvinar is involved with multi-sensory, associative, and higher order cognitive function (*3, 80*). The overall biased distribution of EN1 and EN2 neurons along the medial-to-lateral axis is consistent with the embryonic mouse thalamus (*5*). Epithalamus, which is a neighboring diencephalic region derived from prosomere 2 like the thalamus, contains glutamatergic neurons associated with EN1 signatures **(Fig. 3E)**. This is in line with previously reported similarities in gene expression profiles between epithalamus and thalamus (*29, 81*). However, other neighboring regions, including the subthalamic nucleus and midbrain, contained glutamatergic neurons that clustered separately from EN1 and EN2 (EN-STN, EN-MB) **(Fig. 3E)**.

We next analyzed the distribution of GABAergic neuron subtypes revealed by our MERFISH data (**Fig. 4A; figs. S6A and S7C**). Midbrain-derived *OTX2/SOX14*-positive, *NKX2*.*2*-negative cells (IN1) represented the most abundant population of GABAergic neurons in the thalamus, in line with recent descriptions in the mouse and marmoset (*53*) (**Fig. 4, B and D; figs. S6B and S7C**). In humans, these cells are distributed throughout the thalamus (**Fig. 4, B and D; figs. S6B and S7C**), unlike in mice where they are enriched in the LGN (PMID: **33522480**). GABAergic neurons derived from the prethalamus (IN4) and telencephalon-derived neurons (IN2, IN5) are enriched in the reticular nucleus and zona incerta, but also found sparsely distributed across the thalamic nuclei (**Fig. 4, B and D; fig. S7C**). Abundant IN2 population is also present in the MGE and basal ganglia that were visible in our sections (i.e. caudate, globus pallidus), consistent with their telencephalic identity **(Fig. 4B)**. IN5 is enriched in epithalamus (**fig. S7)**. Finally, *FOXG1*-positive IN6 cluster was observed in the caudate but not the thalamus, suggesting that this sparse population may have been accidentally included in the sc/snRNASeq data generation due to inaccurate dissection **(Fig. 4B)**. We also could not detect IN3, which we hypothesized were pretectal GABAergic neurons, likely due to lack of representation of pretectum in our spatial transcriptomics datasets.

We sought to validate the existence of telencephalon-derived cells in the thalamus by performing immunostaining against FOXG1 in a biologically-independent GW19 sample. Nissl staining revealed the proposed migratory stream between the MGE and the thalamus previously observed in humans, the corpus gangliothalamicus, which is defined as a thin stream of cells starting from the stria terminalis and lining the dorsal edge of the thalamus (**Fig. 4D)**. We observed that cells within this migratory stream stained positive for FOXG1, confirming telencephalic identity **(Fig. 4, F and G)**. Whereas previous reports suggest that these cells migrate predominantly to mediodorsal and pulvinar nuclei, we observe FOXG1+ cells sparsely throughout the thalamus, with enrichment in the reticular nucleus **(Fig. 4, F and G; fig. S11)**. We additionally performed in situ hybridization against *FOXG1* and *CRABP1* across seven sections total collected from three biologically independent thalamus samples. We further confirmed the presence of *FOXG1/CRABP1-*expressing cells within the thalamus and epithalamus, and enrichment in the reticular nucleus, confirming our MERSCOPE results **(fig. S11)**.

We finally assessed the distribution of non-neuronal cell types (**Fig. 5B and fig. S7D)**. LGR6+ astrocytes (AC1) appeared to be the most abundant glial population in lateral sections whereas the bipotent glial progenitors (GL) were the most abundant in medial sections (**Fig. 5, B-D)**, consistent with the lateral-to-medial maturation gradient that was previously observed in the context of neurogenesis in the rodent thalamus (*58, 82*). *SPARCL1+* AC2 astrocytes were enriched in extrathalamic regions such as the midbrain and reticular nucleus, but were also present to a lesser degree in the thalamus (**Fig. 5, B and C; fig. S7D)**. FOXJ1+ ependymal cells are enriched along the periphery of the thalamus and within the centromedian nucleus (**Fig. 5, B and C; fig. S7D)**. Other cell types, including dividing cells, OPCs, oligodendrocytes, microglia, pericytes, fibroblasts were found distributed broadly throughout the thalamus (**Fig. 5, B-D)**.

We identified several genes enriched in anatomically-defined thalamic nuclei. For example, *INHBA, TLL1, ITGA8*, and *KCTD8* were enriched in centromedian nucleus (**fig. S14**). Ventral anterior nucleus was highly enriched for *PENK, GABRG1, TSHZ2*, and *CDH9* (**fig. S14**). Reticular nucleus showed enriched expression of *CNTNAP4, EPHA5, PDE1C*, and *PVALB* (**fig. S14**). Zona incerta was highly enriched for *LHX5, PAX6, ISL1*, and *DLX2* (**fig. S14**). Given the spatial bias we observed for many neuron-enriched genes, we sought to determine whether thalamic nuclei can be molecularly distinguished in midgestation in the human brain. We designed a new gene panel containing 140 genes based on top markers for neuronal cell types, and performed MERSCOPE on a sagittal section from GW21 human thalamus. Clustering analysis revealed segmentation of the thalamus into regions reminiscent of thalamic nuclei independently identified by Nissl staining (**fig. S14**). These results confirm that spatial variation of neuronal gene expression parcellates the thalamus into molecularly-defined nuclei as early as the second trimester.

Together, our study identifies the molecular trajectories of neuronal differentiation in the developing human diencephalon and suggests three principles of neuronal differentiation and spatial organization. One, human thalamic neuronal subtypes begin to emerge during the first trimester of development and start to organize into spatially distinct nuclei during the second trimester. Two, molecularly defined subtypes of glutamatergic neurons are shared across thalamic nuclei, and their composition can be used to distinguish first-order and higher-order thalamic nuclei. Three, we detect six molecular subtypes of GABAergic neurons that contribute to the developing diencephalon in humans. These include prethalamus-derived, telencephalon-derived, and midbrain-derived GABAergic neurons that together contribute substantially to the cellular composition of diencephalic subregions in humans. In particular, we find that midbrain- and telencephalon-derived GABAergic neurons appear in the developing thalamus as early as the second trimester of development.

Our study confirms previously hypothesized migration of telencephalon-derived GABAergic neurons into the thalamus, which may contribute to the expansion of this structure in humans (*54, 55*). These cells maintain expression of *FOXG1* during the second trimester, and form a thin stream along the dorsal edge of the thalamus that coincides with the corpus gangliothalamicus, a migratory pathway between the MGE and the thalamus that was previously identified using Golgi and Nissl stains (*54, 55, 83*). In addition to the previously identified localization of these cells in mediodorsal and pulvinar nuclei, our results indicate that FOXG1-positive cells appear to migrate throughout the thalamus, with particular enrichment in the reticular nucleus. We also observe their presence in the epithalamus, suggesting a greater caudal dispersion of these cells across prosomere 2 of the diencephalon than previously appreciated. Finally, a subset of *FOXG1*-expressing cells expresses *CRABP1*, a gene involved in the retinoic acid signaling pathway. *CRABP1*-positive GABAergic neurons have been recently identified in studies of primate forebrain interneuron diversity (*28, 32*). Consistently, *CRABP1*-positive GABAergic neurons expressed *LHX6* and *MAF* in our single cell sequencing data, suggesting medial ganglionic eminence origin, but did not express *TAC3*. Together, our findings underscore the complexity of neurodevelopmental cell lineage relationships in primates and provide a novel cellular substrate that could contribute to the evolution of the human thalamus.

## Acknowledgements

We thank Matthew Schmitz and members of the Nowakowski lab for providing helpful feedback during manuscript preparation. Widefield imaging data for this study were acquired at the Center for Advanced Light Microscopy - Weill Innovation Center Microscopes. This publication was supported by and coordinated through BICCN. This publication is part of the Human Cell Atlas-www.humancellatlas.org/publications/

## Author contributions

Conceptualization: CNK, DS, TJN

Methodology: CNK, DS, AW, TJN

Investigation: CNK, DS, TJN

Visualization: CNK, DS, TJN

Funding acquisition: TJN

Project administration: TJN Supervision: TJN

Writing – original draft: CNK, DS, TJN

Writing – review & editing: CNK, DS, TJN

## Data and materials availability

Processed scRNA-seq are available on UCSC cell browser at http://cells.ucsc.edu/?ds=dev-thal. Processed MERFISH data are available on Dryad: https://doi.org/10.5061/dryad.98sf7m0q1

## Supplementary Materials

Materials and Methods

Supplementary Text

Figs. S1 to S14

Tables S1 to S3

