## Supplemental Files for "Spatiotemporal molecular dynamics of the developing human thalamus"

**This PDF file includes:**

Materials and Methods
Supplementary Text
Figs. S1 to S14
Tables S1 to S3

### Materials and Methods

Singe cell data processing

scRNA-seq FASTQs were aligned and quantified with STARSolo 2.7.10a with specific parameters for multimapping with expectation-maximization, intron-inclusion, cell barcode and UMI correction: `--soloCBmatchWLtype 1MM_multi_Nbase_pseudocounts --soloUMIdedup 1MM_CR --soloFeatures GeneFull_Ex50pAS Velocyto --soloMultiMappers EM --clipAdapterType CellRanger4 --soloUMIfiltering MultiGeneUMI_CR`. Reference genome annotation used is a modified annotation provided by 10x Genomics (2020-A) based on removing and modifying ambiguous transcripts. The annotations are provided at https://storage.googleapis.com/generecovery/human_GRCh38_optimized_v1_1_velocyto_exonic.gtf.gz. For calling cells, we initially filtered for 500 genes expressed per cell followed by DropletQC thresholding for empty droplets based on the spliced and unspliced ratio from STARSolo Velocyto's quantification. DropletQC rescue parameters were set at 0.1 for nf_rescue and 1000 for umi_rescue. The DropletQC filtered matrix was then removed for ambient RNA with FastCAR with recommended cutoff and contaminationChanceCutoff set to 0.05 and stop iteration at 200. The ambient RNA removed matrix was then used to compute doublet prediction with scds' hybrid mode. The threshold for scds hybrid score was stepwise cutoffs based on cells called with more stringent doublet filtering with higher number of cells called. Mitochondrial threshold was the final step for filtering at 20% maximum mitochondrial content.

Single cell analysis

Seurat's SCTransform v2 workflow was used for normalization set to 2000 most variable genes followed by filtering for mitochondrial, ribosomal genes, and MALAT1. The residuals were input into PCA and batch corrected with harmony over 20 principal components for the first trimester and 30 for the second trimester with theta=2 over the sample metadata. UMAP dimensional reduction was done on the harmony corrected components with min.dist=0.3. De novo clustering was done with Louvain with multilevel refinement with the resolution set 0.5 for the first trimester and 0.4 for the second trimester. TF activity was computed using Dorothea and pathway activity with progeny. Visualization tools used include dittoSeq, SCpubr, and scCustomize. All analysis scripts are available at https://github.com/cnk113/thalamus-analysis.

MERSCOPE panel design, tissue preparation, and imaging

We selected a panel of 140 genes to perform MERSCOPE spatial transcriptomics profiling using Vizgen's commercial platform. Genes were selected based on cluster markers identified from sc/snRNA-seq data, including canonical and novel markers against neuronal, glial, and non-neural cell types in the first and second trimester thalamus. The full gene panel is available in Supplemental Table 3. A second panel of 140 genes was designed to focus on markers associated with neuronal clusters in the second trimester thalamus for the experiment described in fig. S14, and is also available in Supplemental Table 3.

The panel was used to stain 10 μm thick sections generated from thalamus tissue (GW10, GW16, GW21, GW23). Thalamus tissue was dissected from an intact hemisphere on the day of collection, flash frozen in a liquid nitrogen-chilled isopentane bath, and then immediately frozen in ice-cold OCT while documenting its orientation relative to the rest of the brain. The tissue was serially sectioned in the sagittal orientation on a Leica CM1860 using a specimen head temperature of -15°C. One 100 μm thick section was used for generating RNA for confirming a

RIN score greater than seven as a quality control measurement. RNA was isolated using the Direct-Zol RNA purification kit (Zymo) according to the manufacturer's recommendations. RIN scores were measured using the Agilent RNA 6000 Pico Kit. The mediolateral position of subsequent sections were gauged by performing Nissl stain using the Neurotrace 500/525 dye (ThermoFisher) and comparing the cytoarchitecture with the Bayer and Altman Reference Atlas. Once we identified a suitable region of the thalamus for spatial profiling, we collected two sections on Superfrost Plus slides, one for Nissl staining and one for RNAscope, and collected a subsequent section for MERSCOPE on a glass coverslip provided by Vizgen.

Tissue preparation and imaging was performed following Vizgen's recommended procedures for fresh frozen tissue. The MERSCOPE slide with the tissue mounted was fixed using 4% PFA for 15 minutes, washed three times in PBS, and permeabilized overnight in 70% ethanol at 4°C. The next day, encoding probe hybridization was performed for 36-48 hours, followed by gel embedding and clearing for 24-96 hours following the non-resistant fresh frozen tissue guidelines. Finally, the tissue was stained using the DAPI and PolyT Staining Reagent provided by the kit for fifteen minutes. The MERSCOPE instrument was configured and the tissue was imaged according to the Vizgen MERSCOPE Instrument User Guide.

Immunohistochemistry and fluorescent in situ hybridization

Fluorescent in situ hybridization was performed using the RNAscope Multiplex Fluorescent v2 Assay against human *FOXG1* (probe ID: 420991-C3) and *CRABP1* (probe ID: 855321-C2) according to the manufacturer's protocol for fresh frozen tissue. Immunohistochemistry was performed on 10 µm thick cryosections generated from GW19 thalamus tissue that had been fixed in 4% PFA overnight, dehydrated in 30% sucrose for 48 hours, OCT-embedded, and stored at -80°C. Antigen retrieval was performed prior to immunostaining by incubating slides for 15 minutes in citrate buffer (pH 6) preheated to 95°C using a pressure cooker. Slides were washed three times in PBS containing 0.1% Tween-20, and simultaneously permeabilized and blocked using PBS containing 5% BSA and 0.3% Triton-X. Slides were stained overnight at 4°C in blocking buffer containing antibodies against FOXG1 (Abcam 196868, 1:500 dilution) and TCF7L2 (Millipore 05-511 clone 6H5-3, 1:250 dilution). The next day, slides were washed three times for 10 minutes each in PBS containing 0.1% Tween-20, and incubated in blocking buffer containing secondary antibodies (ThermoFisher A32766 and A32795, each diluted 1:1000) for one hour. Slides were washed three times in PBS, stained with DAPI (ThermoFisher 62247), and mounted using Prolong Gold (Invitrogen, P36930) and #1.5 coverslips (Azer Scientific, 1152460).

Tilescan images were collected using the Leica Widefield microscope equipped with a Hamamatsu Flash 4.0 camera using either a 20X air objective for RNAscope or a 10X air objective for immunostaining. High magnification images were collected using a Leica SP8 confocal microscope with a 63X oil objective.

Spatial transcriptomics analysis

Cell segmentation was performed using the polyT and DAPI stains. The output of Vizgen's processing software with DAPI segmentation was used for input into Seurat's SCTransform v2 normalization. The matrix was initially filtered for a minimum of 20 counts and a maximum of 100 unique genes expressed per cell. Label transfer from second trimester to spatial data was done using TransferAnchors and MapQuery on the harmony reductions with cluster labels. All analysis scripts are available at https://github.com/cnk113/thalamus-analysis.

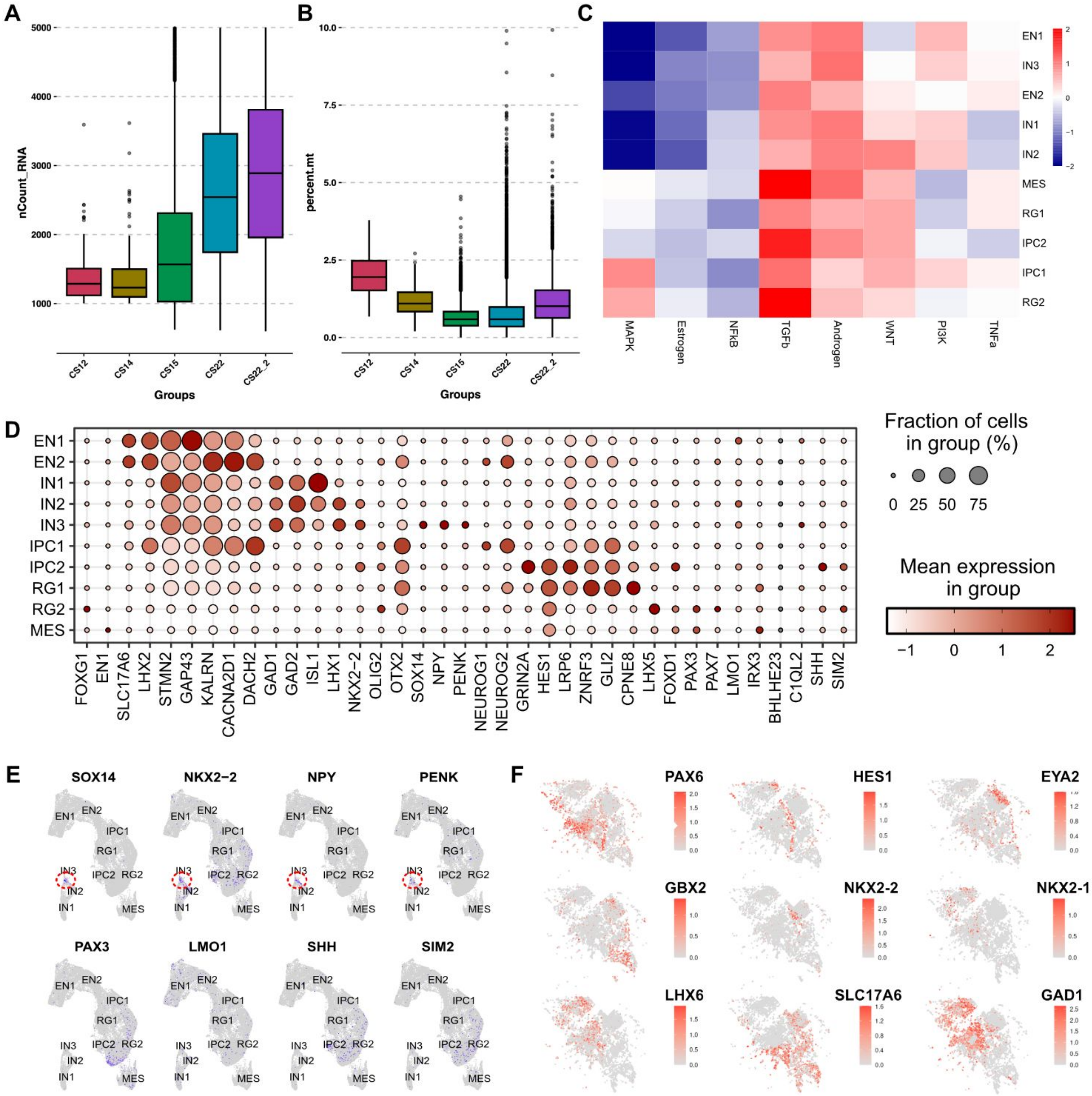


**Fig. S1. A.** UMI per cell across samples in the first trimester scRNA-seq dataset. **B.** Percent mitochondria content across samples. **C.** Pathway activity across cell states in the first trimester. **D.** Dotplot of additional marker genes across cell states in the first trimester. **E.** Feature plots of log-normalized gene expression projected across cells in UMAP. **F.** Spatial feature plots of log-normalized gene expression of cell type markers across GW10 MERFISH spatial coordinates.

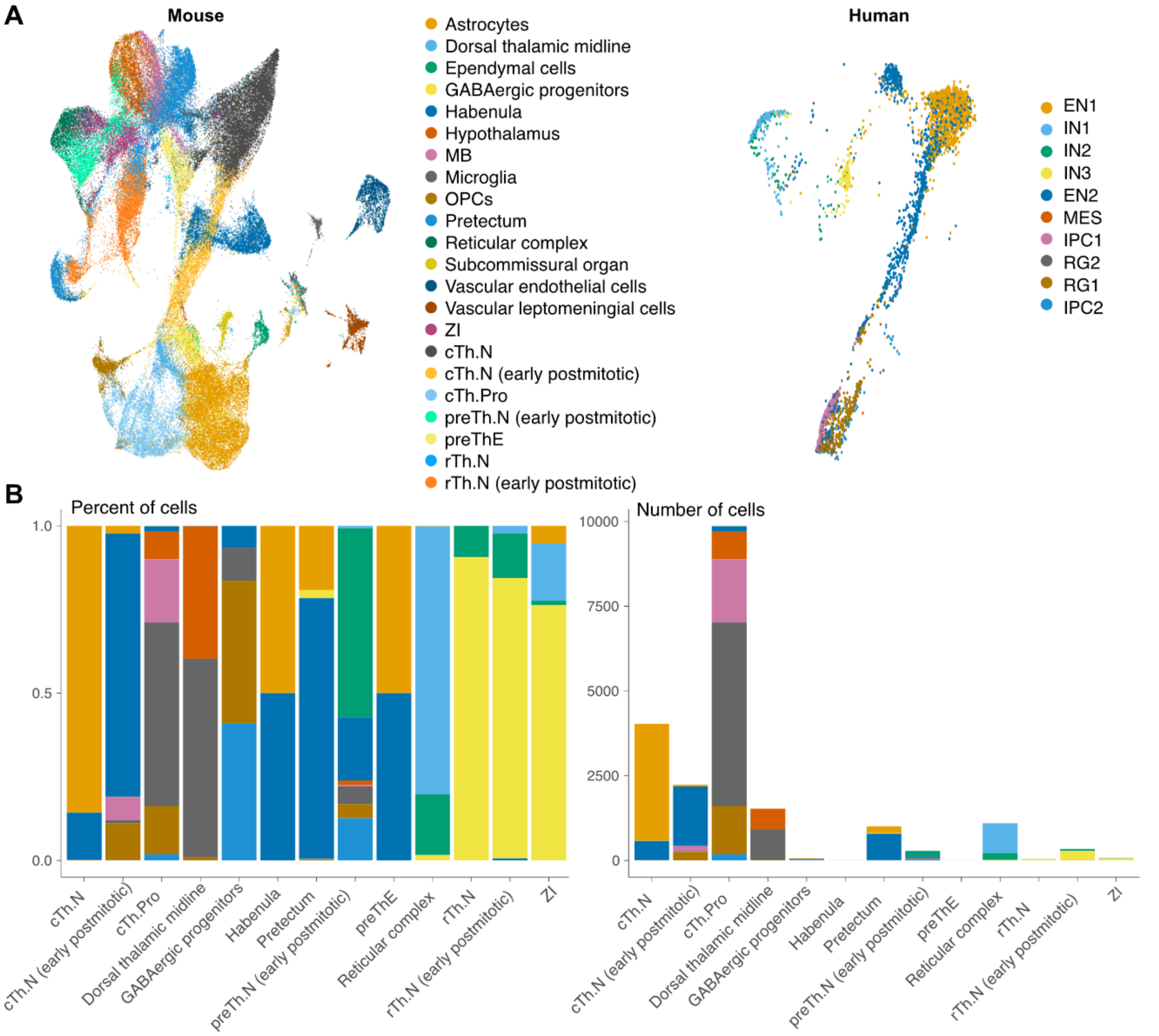


**Fig. S2. A.** UMAP embedding of the mouse reference atlas (left) and query cells from the first trimester thalamus (right). **B.** Proportion of cells from the first trimester stratified across mouse annotations (left) and raw cell numbers (right).

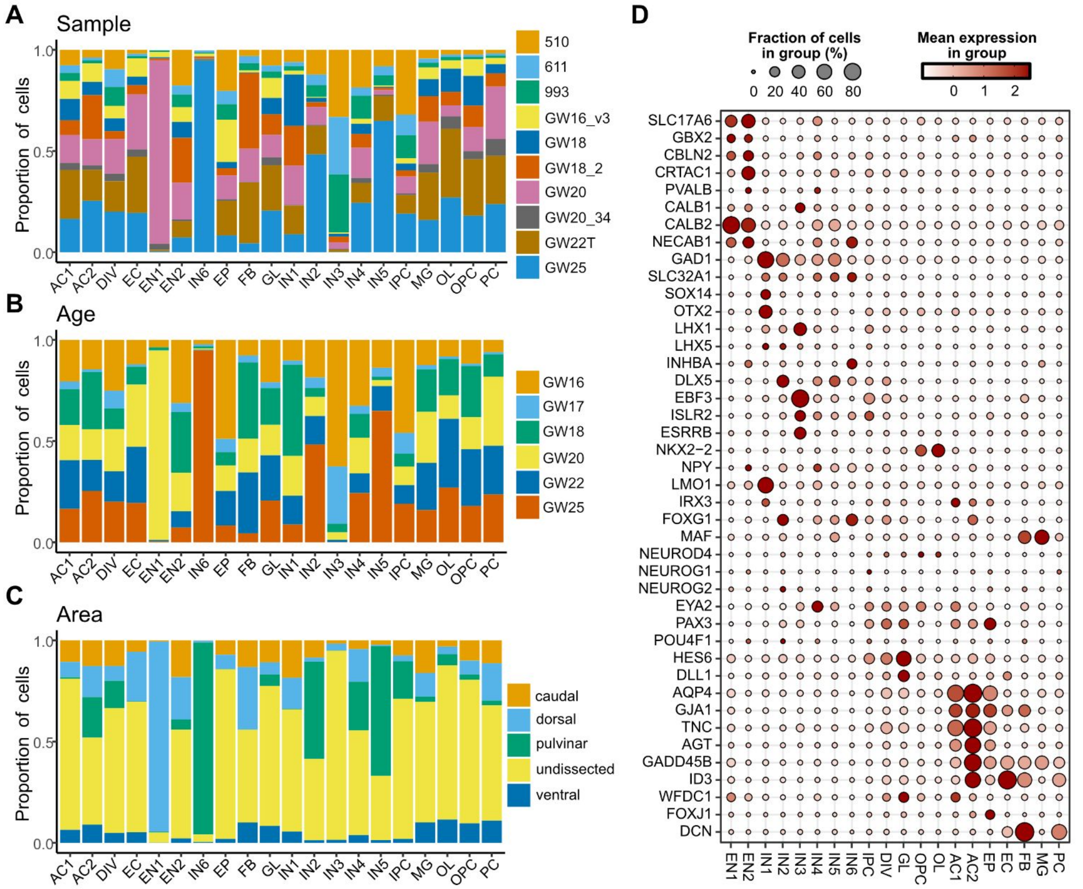


**Fig. S3. A.** Barplot across cell states in the second trimester filled by sample, **B.** age, and **C.** area. **D.** Dotplot depicts expression of marker genes.

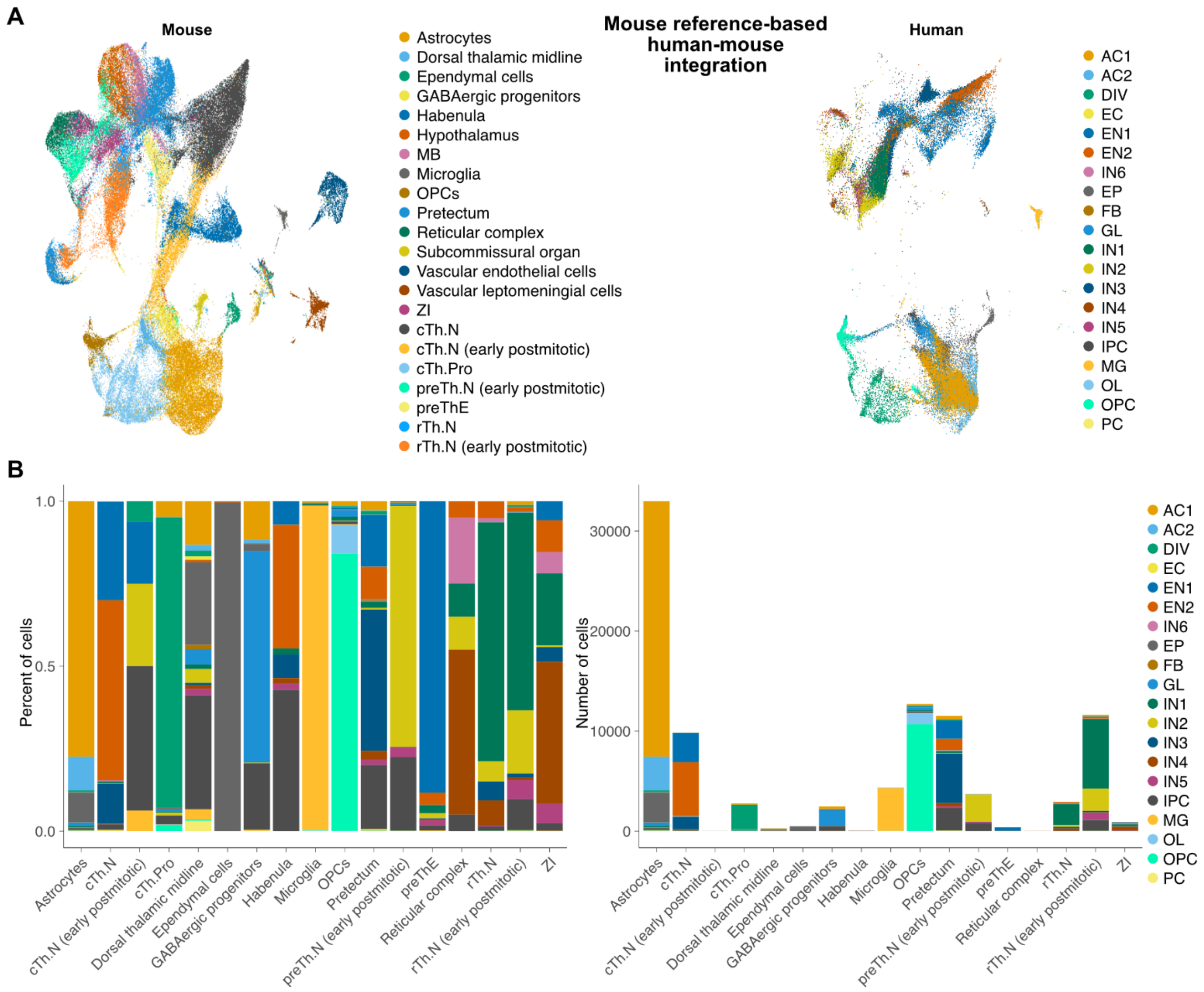


**Fig S4. Evolutionary divergence of inhibitory neurons between human and mouse. A.** UMAP embedding of mouse E14.5 - E18.5 developmental thalamus atlas onto second trimester embedding. **B.** Cell count of human second trimester thalamus atlas across cell cluster labels. **C.** (Left) Relative proportion of mouse scRNA-seq profiles across predicted mapping from human second trimester labels. (Right) Raw count of proportion from developing mouse thalamus across second trimester labels. Mouse labels: cTh: caudal thalamus, rTh: rostral thalamus, preTh: pre thalamus, MB: midbrain, and ZI: zona incerta.

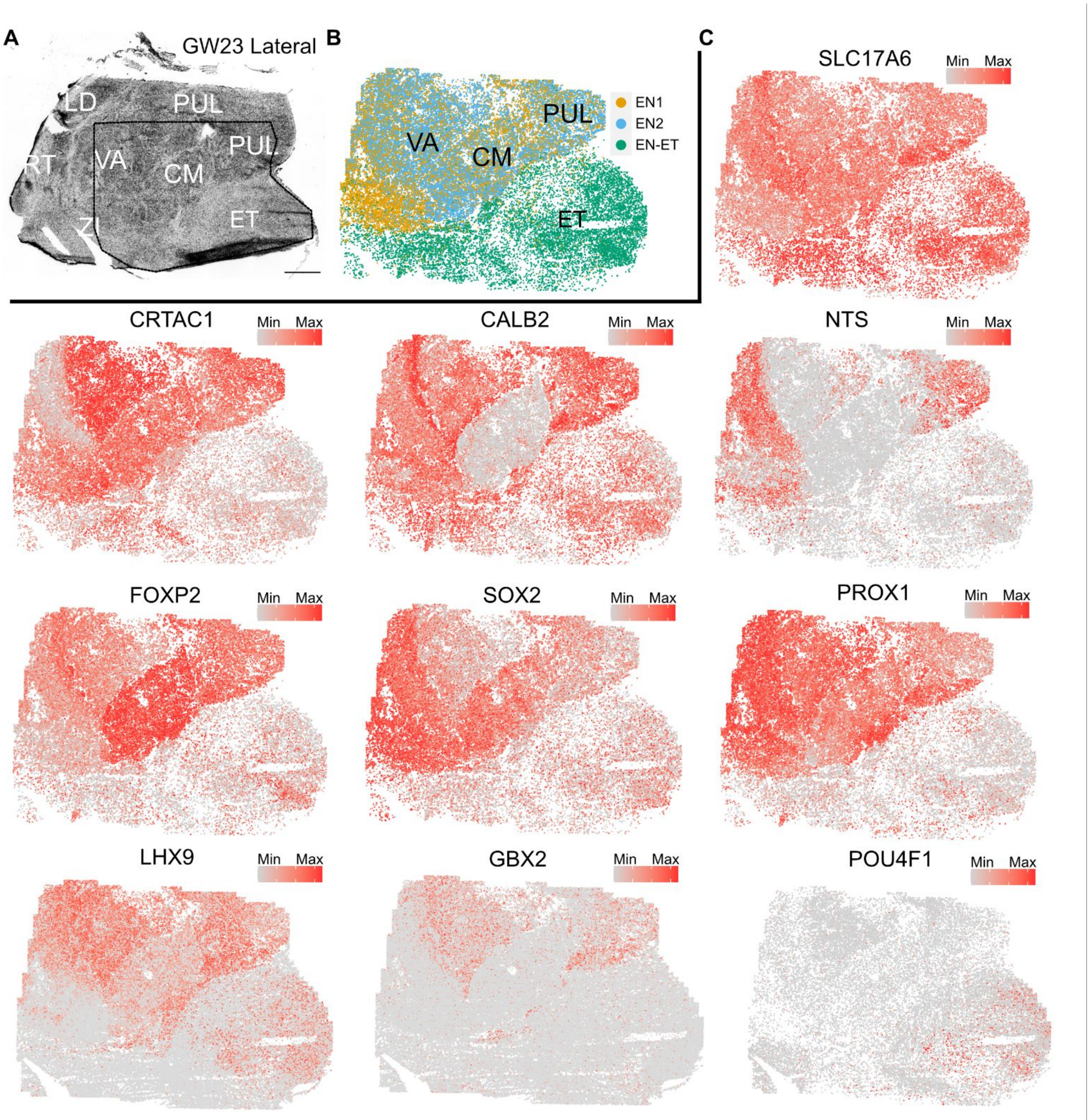


**Fig S5. A.** Nissl stain of a sagittal section collected from a lateral region of GW23 human thalamus. **B.** Spatial distribution of glutamatergic neuron subtypes EN1 and EN2, which correspond to clusters from scRNA-seq, and EN-ET, which correspond to glutamatergic neurons in the epithalamus. **C.** Spatial feature plots projecting log normalized expression of glutamatergic neuron marker genes within the glutamatergic neuron subset of the MERFISH dataset.

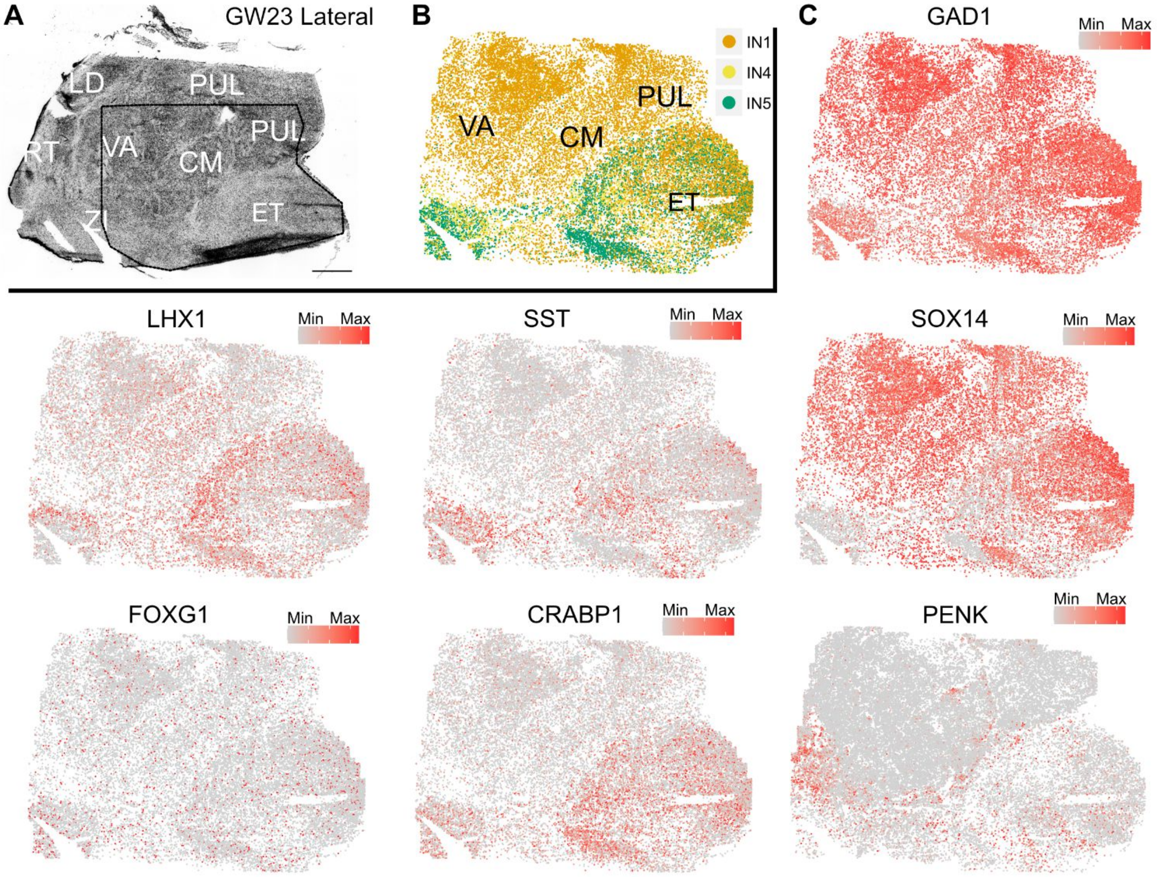


**Fig S6. A.** Nissl stain of a sagittal section collected from a lateral region of GW23 human thalamus. **B.** Spatial distribution of detected GABAergic neuron subtypes IN1, IN4, and IN5, which correspond to what are likely midbrain-derived, prosomere 3-derived, and telencephalon-derived populations. **C.** Spatial feature plots projecting log normalized expression of glutamatergic neuron marker genes within the glutamatergic neuron subset of the MERFISH dataset.

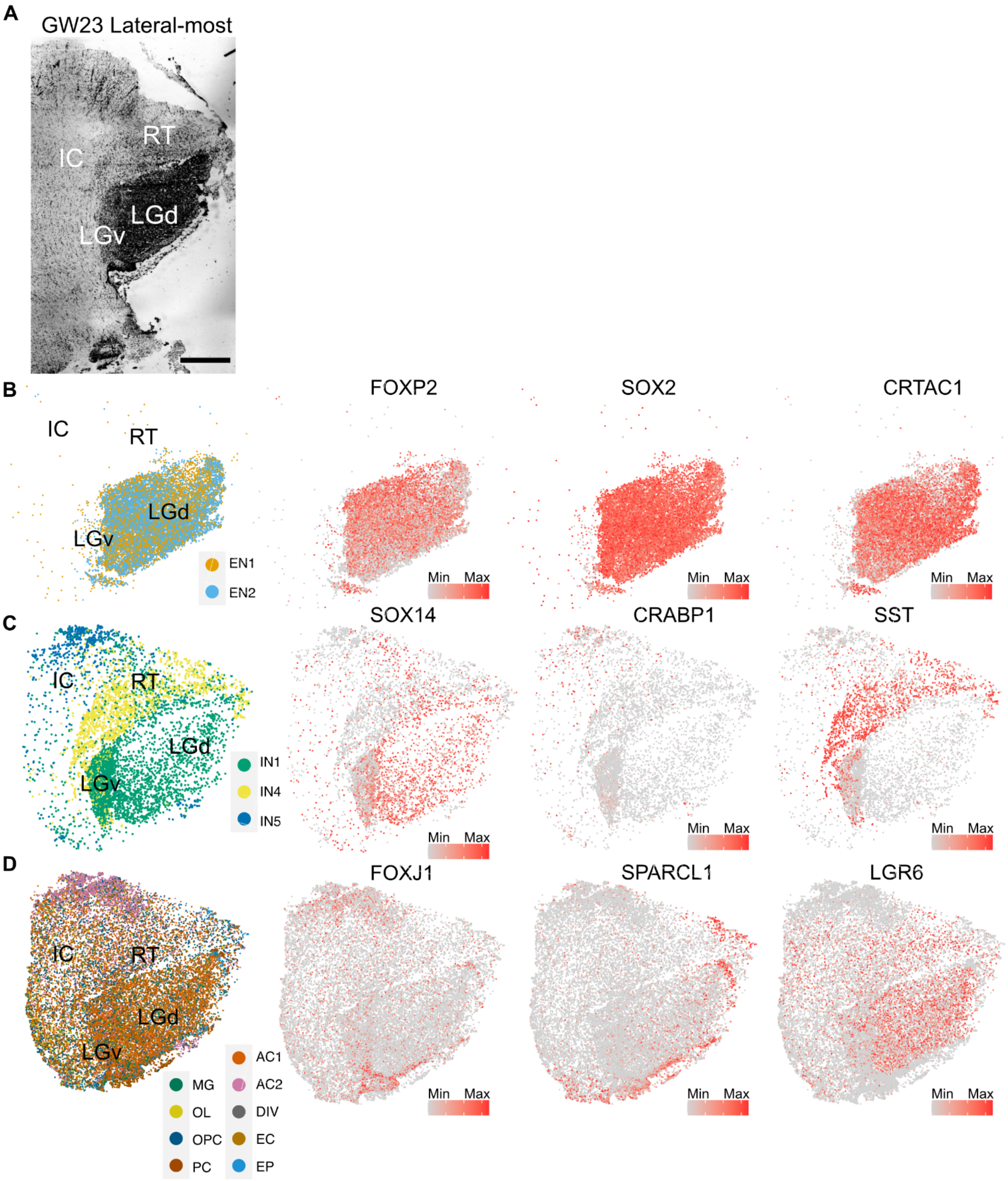


**Fig S7. A.** Nissl stain of a sagittal section collected from the lateral end of GW23 human thalamus. **B.** Spatial distribution (left) and feature marker gene expression (right) of glutamatergic neuron subtypes, **C.** GABAergic neuron subtypes, and **D.** non-neural subtypes.

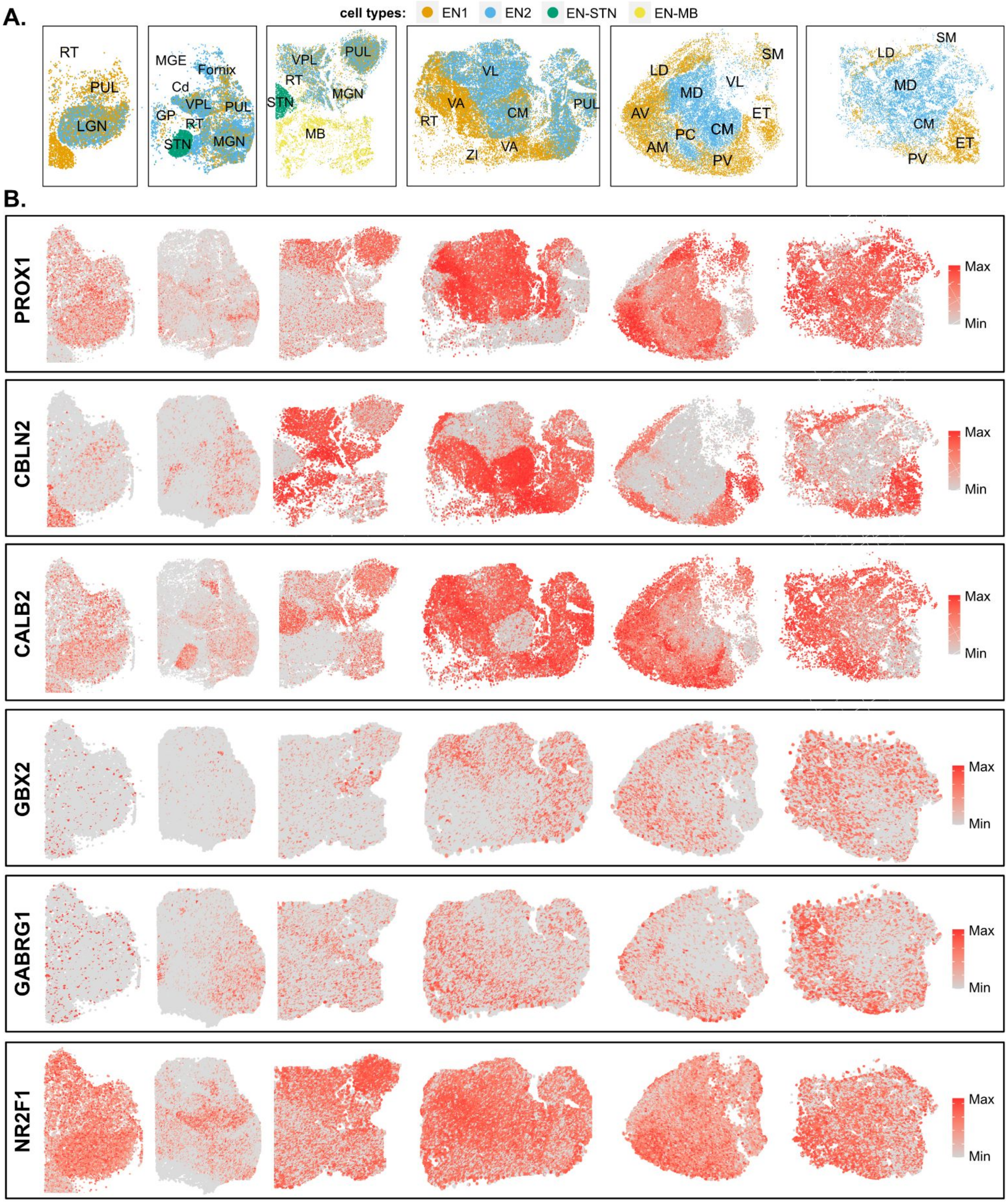


**Fig. S8. A.** Spatial distribution of glutamatergic neuron subtypes. **B.** Spatial feature plots of markers for glutamatergic neuron subtype EN1.

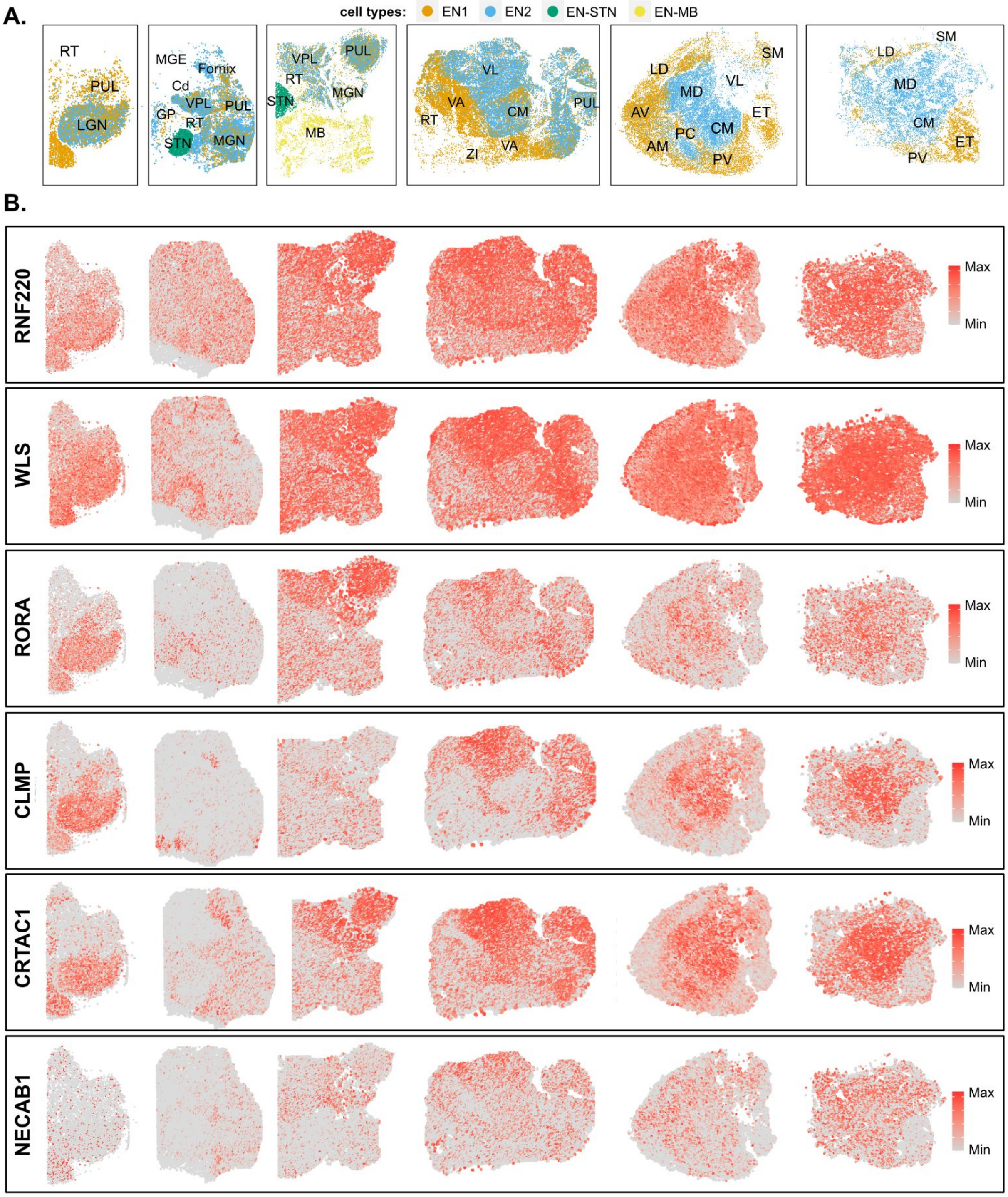


**Fig. S9. A.** Spatial distribution of glutamatergic neuron subtypes. **B.** Spatial feature plots of markers for glutamatergic neuron subtype EN2.

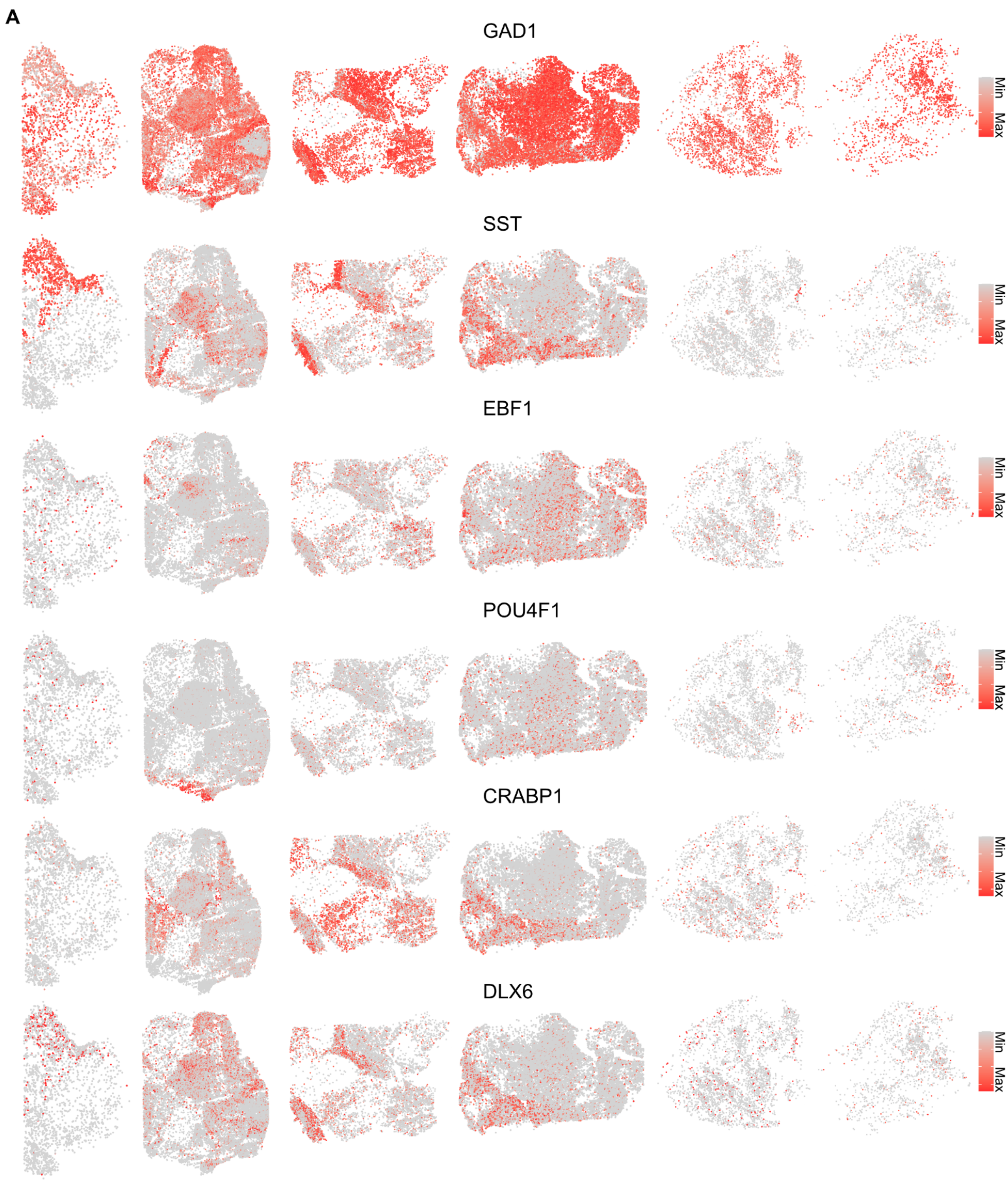


**Fig. S10. A.** Spatial feature plots of GABAergic neuron subtype marker genes within datasets subsetted for GABAergic neurons.

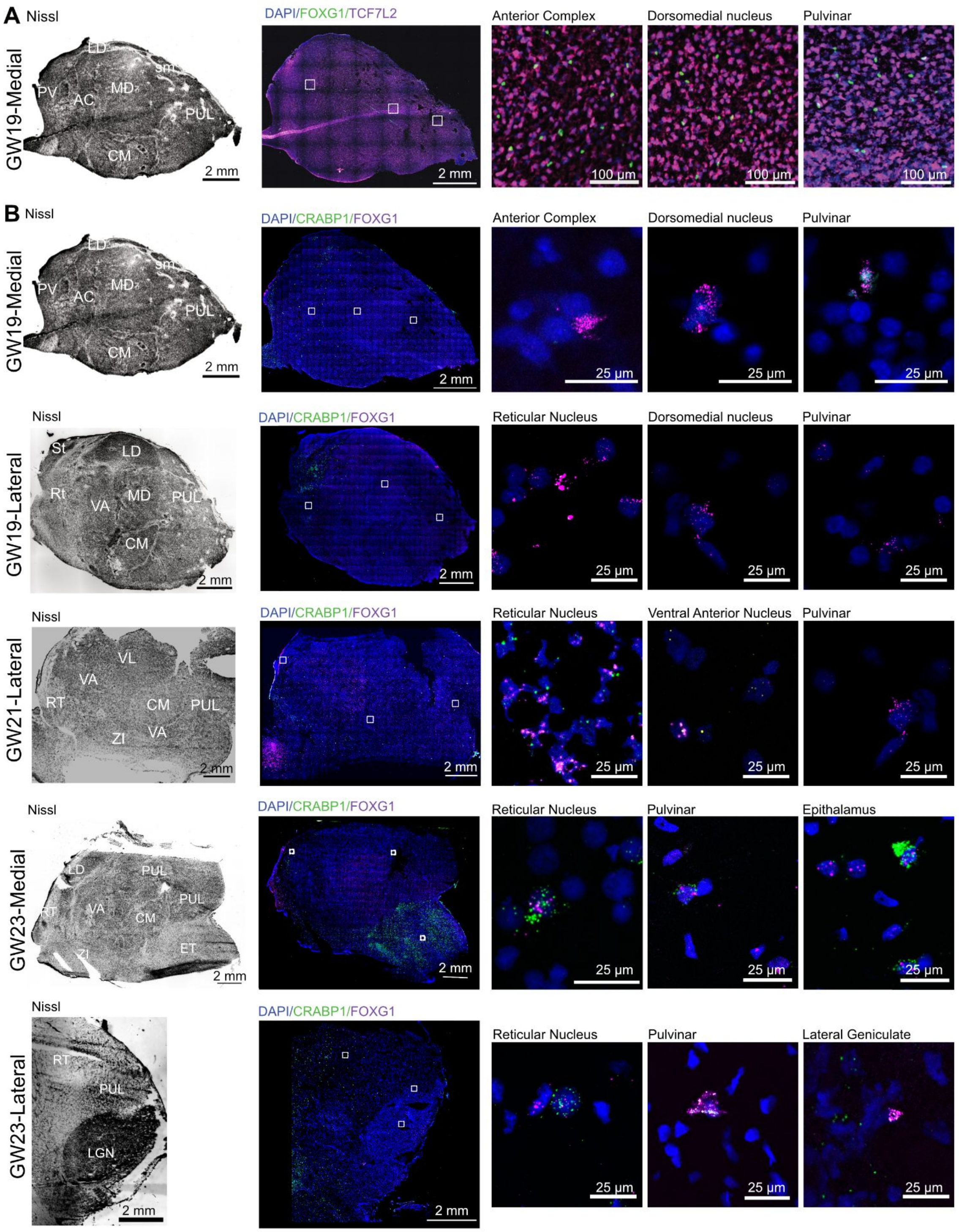


**Fig. S11. A.** Nissl stain (left) and immunohistochemistry (right) against FOXG1 and TCF7L2 in a sagittal section collected from the medial side of GW19 human thalamus. Nissl and immunostains were performed on adjacent sections. Higher magnification images were collected using a 20X objective to visualize regions indicated by white insets in the full image. **B.** Nissl stains (left) and RNAscope (right) against *CRABP1* and *FOXG1* performed on sagittal sections collected from three biologically independent mid-gestation thalamus samples (GW19, GW21, GW23). Nissl stains and RNAscope experiments were conducted on adjacent sections. High magnification images were collected using a 63X objective to visualize regions indicated by white insets in the full image. **Abbreviations:** RT - Reticular Nucleus, PUL - Pulvinar, LGN - Lateral Geniculate Nucleus, MGE - Medial Ganglionic Eminence, Cd - Caudate, GP - Globus Pallidus, STN - Subthalamic Nucleus, VPL - Ventral Posterior Lateral Nucleus, MGN - Medial Geniculate Nucleus, MB - Midbrain, VA - Ventral Anterior Nucleus, VL - Ventral Lateral Nucleus, CM - Centromedian Nucleus, ZI - Zona Incerta, AV - Anteroventral Nucleus, AM - Anteromedial Nucleus, LD - Dorsolateral Nucleus, MD - Dorsomedial Nucleus, PC - Paracentral Nucleus, PV - Paraventricular Nucleus, SM - Stria Medullaris, ET - Epithalamus.

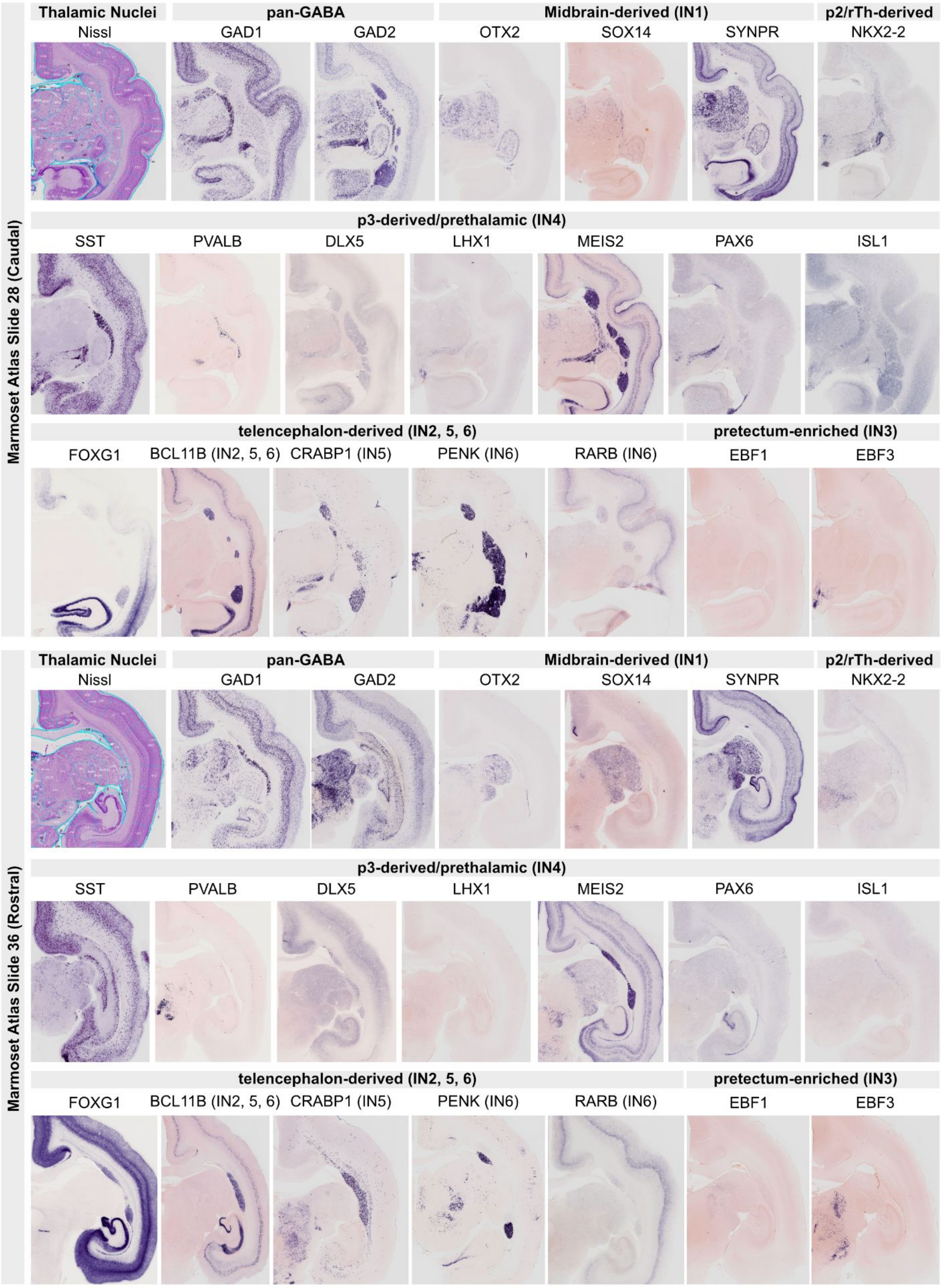


**Fig. S12.** In situ hybridizations from marmoset gene atlas (https://gene-atlas.brainminds.jp/) visualizing genes relating to GABAergic neuron subtypes in second trimester scRNA-seq dataset

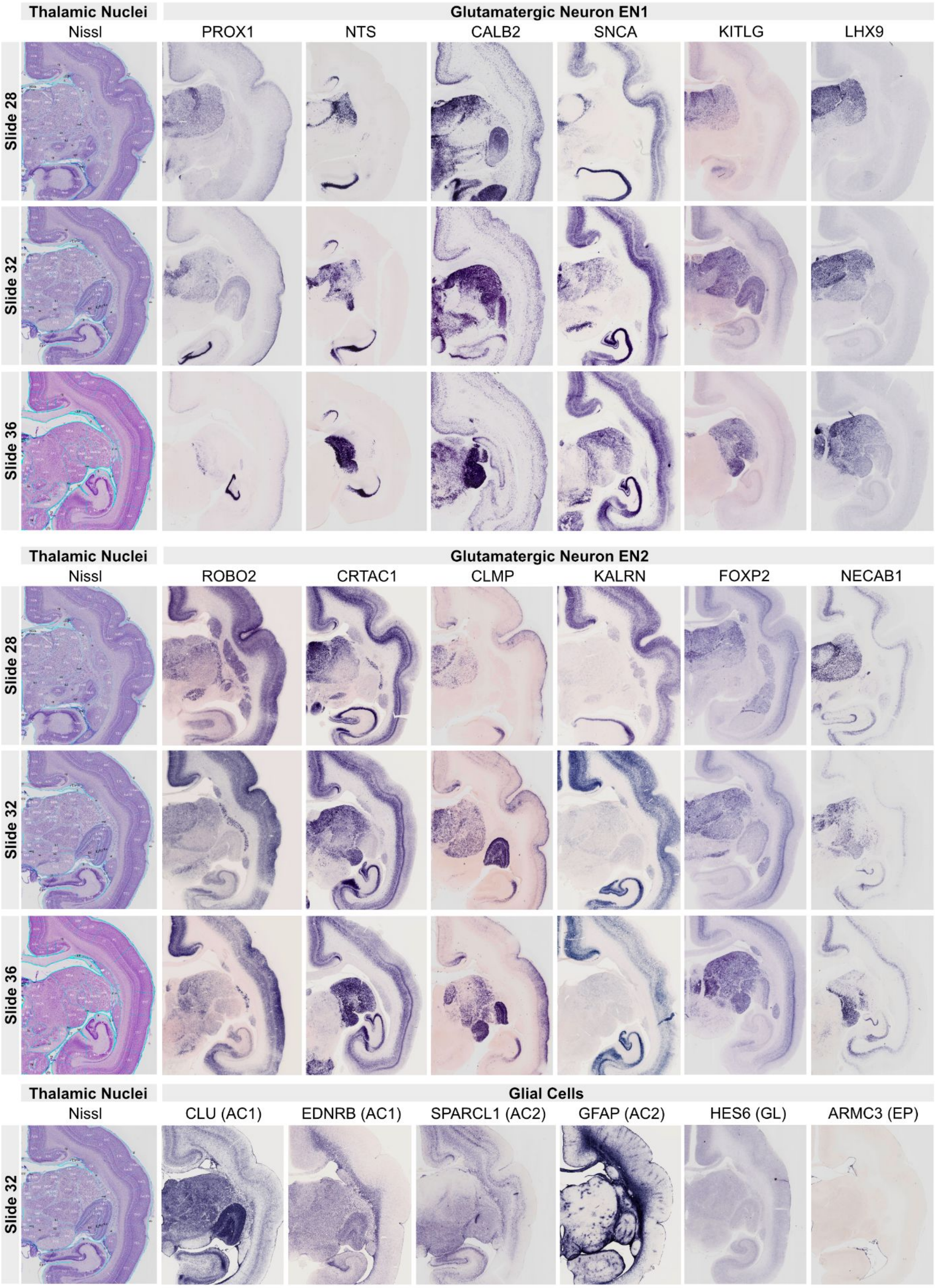


**Fig. S13.** In situ hybridizations from marmoset gene atlas (https://gene-atlas.brainminds.jp/) visualizing genes relating to glutamatergic neuron and glial subtypes in second trimester scRNA-seq dataset

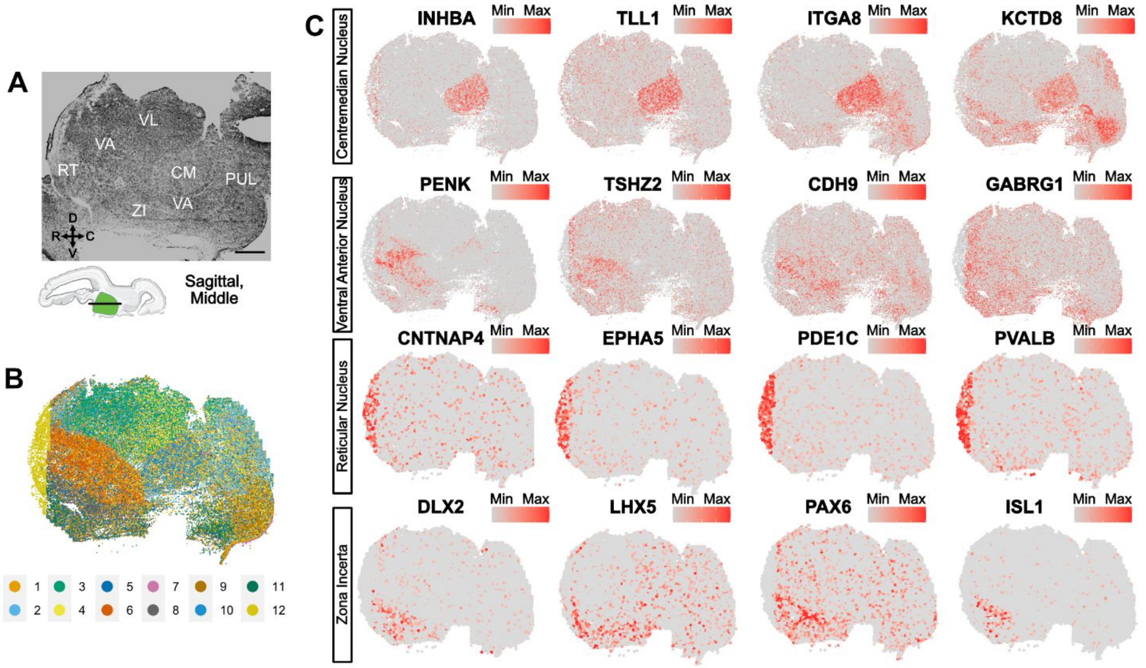


**Fig. S14. A.** We performed MERFISH spatial transcriptomics using a 140 gene panel composed of neuronal subtype markers on a sagittal section from GW21 human thalamus that was adjacent to the GW21 section in the main text. Nissl stain is featured here. **B.** Spatial clustering analysis reveals partitions in the thalamus reminiscent of thalamic nuclei revealed by Nissl stain. **C.** Spatial feature plots for genes enriched for centromedian nucleus, ventral anterior nucleus, reticular nucleus, and zona incerta.

**Table S1. Marker genes for first trimester cell states.** Differentially expressed genes were identified using the wilcoxon rank sum test on SCTransform normalized counts comparing clusters in the first trimester thalamus scRNA-seq dataset.

**Table S2. Marker genes for second trimester cell states.** Differentially expressed genes were identified using the wilcoxon rank sum test on SCTransform normalized counts comparing clusters in the second trimester thalamus scRNA-seq dataset.

**Table S3. MERFISH gene panel.** Genes used for the MERFISH probe panel.
